## Supplementary material for "Sex and Power: sexual dimorphism in trait variability and its eco-evolutionary and statistical implications": FullCode&Supplemental_Information


Code 

- Show All Code
- Hide All Code

### Sex and Power: Sexual dimorphism in trait variability and its eco-evolutionary and statistical implications

##### Electronic Supplementary Material

###### Susanne Zajitschek, Felix Zajitschek, Russell Bonduriansky, Robert Brooks, Will Cornwell, Daniel Falster, Malgorzata Lagisz, Jeremy Mason, Alistair Senior, Daniel Noble, & Shinichi Nakagawa

#### 2020-09-12

### 1 General Introduction

This supplementary material follows the general structure described in our methods and is outlined here in Figure 1.1. Generally speaking, the first few sections describe the processes by which the data were cleaned and re-organised for analysis. This is followed by how the first order and second order meta-analyses were conducted. Supplemental tables and figures that are referenced within the main manuscript can be found at the end of the document. Various parts of the workflow can be conveniently referenced using the table of contents panel (located along the side). We will refer back to this figure as we describe the various aspects of the code and data processing in relevant places.

Figure 1.1: Workflow of data process and meta-analysis

### 2 Set-up

#### 2.1 Packages

```
library(pacman)
pacman::p_load(readr, dplyr, metafor, devtools, purrr, tidyverse, tidyr, tibble, 
    kableExtra, robumeta, ggpubr, ggplot2, png, grid, here, pander)
```

#### 2.2 Functions

Functions needed for preparing the data for meta analyses

##### 2.2.1 Loading and cleaning data

These functions will load the raw and and do some cleaning to make it ready for further processing and analysis.

```
# loads the raw data, setting some default types for various columns

load_raw <- function(filename) {
    read_csv(filename, col_types = cols(.default = col_character(), project_id = col_character(), 
        id = col_character(), parameter_id = col_character(), age_in_days = col_integer(), 
        date_of_experiment = col_datetime(format = ""), weight = col_double(), phenotyping_center_id = col_character(), 
        production_center_id = col_character(), weight_date = col_datetime(format = ""), 
        date_of_birth = col_datetime(format = ""), procedure_id = col_character(), 
        pipeline_id = col_character(), biological_sample_id = col_character(), biological_model_id = col_character(), 
        weight_days_old = col_integer(), datasource_id = col_character(), experiment_id = col_character(), 
        data_point = col_double(), age_in_weeks = col_integer(), `_version_` = col_character()))
}

# Apply some standard cleaning to the data
clean_raw_data <- function(mydata) {
    
    group <- read_csv(here("data", "ParameterGrouping.csv"))
    
    tmp <- mydata %>% 
    # Filter to IMPC source (recommend by Jeremey in email to Susi on 20 Aug 2018)
    filter(datasource_name == "IMPC") %>% 
    # standardise trait names
    mutate(parameter_name = tolower(parameter_name)) %>% 
    # remove extreme ages
    filter(age_in_days > 0 & age_in_days < 500) %>% 
    # remove NAs
    filter(!is.na(data_point)) %>% 
    # subset to reasonable set of variables, date_of_experiment used as an indicator
    # of batch-level effects
    select(production_center, strain_name, strain_accession_id, biological_sample_id, 
        pipeline_stable_id, procedure_group, procedure_name, sex, date_of_experiment, 
        age_in_days, weight, parameter_name, data_point) %>% 
    # sort
    arrange(production_center, biological_sample_id, age_in_days)
    
    # filter to groups with > 1 centre
    merge(tmp, tmp %>% group_by(parameter_name) %>% summarise(center_per_trait = length(unique(production_center, 
        na.rm = TRUE)))) %>% filter(center_per_trait >= 2) %>% 
    # Define population variable
    mutate(population = sprintf("%s-%s", production_center, strain_name)) %>% 
    # add grouping variable: these were decided based on functional groups and
    # procedures
    mutate(parameter_group = group$parameter[match(parameter_name, group$parameter_name)]) %>% 
        
    # Assign unique IDs (per trait) each unique parameter_name (=trait,use trait
    # variable) gets a unique number ('id')
    
    # We add a new variable, where redundant traits are combined [note however, at
    # this stage the dataset still contains nonsensical traits, i.e. traits that may
    # not contain any information on variance]
    mutate(id = match(parameter_name, unique(parameter_name))) %>% as_tibble()
}
```

##### 2.2.2 Sub-setting data

Create a function for sub-setting the data to choose only one data point per individual per trait: “data\_subset\_parameterid\_individual\_by\_age”

```
data_subset_parameterid_individual_by_age <- function(mydata, parameter, age_min = 0, 
    age_center = 100) {
    tmp <- mydata %>% filter(age_in_days >= age_min, id == parameter) %>% # Take results for single individual closest to age_center
    mutate(age_diff = abs(age_center - age_in_days)) %>% group_by(biological_sample_id) %>% 
        filter(age_diff == min(age_diff)) %>% select(-age_diff)
    
    # Still some individuals with multiple records (because same individual appears
    # under different procedures, so filter to one record)
    j <- match(unique(tmp$biological_sample_id), tmp$biological_sample_id)
    tmp[j, ]
}
```

##### 2.2.3 Population statistics

Create a function called: “calculate\_population\_stats”
This function groups animals from the same strain and institution together. This is done for each trait separately, and only for traits that have been measured in both sexes. Any group containing fewer than 6 individuals is excluded.

```
# Function re-organises the data so that the male and female data are side-by-side as columns to make effect size calculations easier.
multi_spread <- function(df, key, value) {
    # quote key
      keyq <- rlang::enquo(key)
    # break value vector into quotes
    valueq <- rlang::enquo(value)
         s <- rlang::quos(!!valueq)
        
         df                           %>% 
        gather(variable, value, !!!s) %>%
        unite(temp, !!keyq, variable) %>%
        spread(temp, value)
}


# Function will calculate population stats and exclude data that is irrelevant and not useful, it will also just convert the data straight away to wide format making it ready for effect size calculations

calculate_population_stats <- function(mydata, min_individuals = 5){
    mydata %>% 
    group_by(population,
             production_center,  
             strain_name,  
             sex) %>%
    summarise(trait  = parameter_name[1],
              x_bar  = mean(data_point, na.rm = TRUE),
              x_sd   =   sd(data_point, na.rm = TRUE),
              n_ind  = n()) %>%
    ungroup() %>%
    filter(n_ind > min_individuals) %>%   
    group_by(population) %>%
    filter(all(c("male", "female") %in% sex)) %>% # Removes any entries where only 1 sex is present
    multi_spread(sex, c(x_bar, x_sd, n_ind))      # Spreads the data out into wide format, so sexes are beside each other which is more typical of meta-analytic data
}
```

##### 2.2.4 Calculate effect sizes and sample variances

Function: “create\_meta\_analysis\_effect\_sizes”

```
# This function takes the data and computes lnCVR, lnVR and ROM (lnRR) using male
# and female data to generate effect size statistics for meta-analyses
create_meta_analysis_effect_sizes <- function(mydata, measure = c("CVR", "VR", "ROM")) {
    for (i in 1:length(measure)) {
        mydata <- metafor::escalc(m1i = male_x_bar, m2i = female_x_bar, sd1i = male_x_sd, 
            sd2i = female_x_sd, n1i = male_n_ind, n2i = female_n_ind, data = mydata, 
            measure = measure[i], var.names = c(paste0("effect_size_", measure[i]), 
                paste0("sample_variance_", measure[i])), append = TRUE)
    }
    return(mydata)
}
```

#### 2.3 Load & clean data

We have already done this step and provide a cleaned up file which is less computationally intensive to deal with. The file has been saved in a folder called `export`. However, the `.csv` is provided in case this is preferred and can be imported and processed using following the steps below: [This code is currently excluded from showing in this `.html` file, but can be viewed and enabled / run in the down-loadable .rmd file. However, it requires the .gz file containing the raw data in the “export” folder. Unfortunately, that file can not be provided via Github, as it is too large (274MB). However, it is freely accessible and uploaded to zenodo.org (https://zenodo.org/deposit/3759701)

```
# Load raw data - save cleaned dataset as RDS for reuse
data_raw <- load_raw(here("data", "dr7.0_all_control_data.csv.gz"))
dir.create("export", F, F)

data <- data_raw %>% clean_raw_data()
saveRDS(data, "export/data_clean.rds")
```

For analysis we load the RDS created above and other datasets

```
data <- readRDS(here("export", "data_clean.rds"))

procedures <- read_csv(here("data", "procedures.csv"))
```

#### 2.4 Mean-variance relationships

We can also have a look at the mean-variance relationships in these data (Figure 2.1). Note that these data have not been filtered to the same extent that the data is that is used in the meta-analyses, but this is simply here to demonstrate that mean-variance relationships exist and are strong providing strong justification for using the log coefficient of variation ratio (lnCVR).

```
# First, clean up the data

 results <- data %>%
            group_by(population,
             production_center,  
             strain_name,  
             parameter_name,
             sex) %>%
    summarise(trait  = parameter_name[1],
              x_bar  = mean(data_point, na.rm = TRUE),
              x_sd   =   sd(data_point, na.rm = TRUE),
              n_ind  = n()) %>%
    ungroup() %>%
    filter(n_ind > 5) %>%   # NOTE here that you are excluding data with 5 exactly....
    group_by(population) %>%
    filter(all(c("male", "female") %in% sex)) %>% # Removes any entries where only 1 sex is present
    multi_spread(sex, c(x_bar, x_sd, n_ind)) %>%
    mutate(lnMean_m = log(male_x_bar),
           logSD_m  = log(male_x_sd),
           lnMean_f = log(female_x_bar),
           lnSD_f   = log(female_x_sd)) %>%
    filter(!is.na(logSD_m) & !is.inf(logSD_m)) %>%
    filter(!is.na(lnSD_f) & !is.inf(lnSD_f))
                 
male <- ggplot(results, aes(x = lnMean_m, y = logSD_m)) +
    geom_point(color = "#66C2A5", size = 2) +
   geom_smooth(method = "lm", se = FALSE, color="black") + 
    labs(x = "log (Mean)",
         y = "log (SD)") + 
    theme_classic()

female <- ggplot(results, aes(x = lnMean_f, y = lnSD_f)) +
    geom_point(color = "#FC8D62", size = 2) +
  geom_smooth(method = "lm", se = FALSE, color="black")+
    labs(x = "log (Mean)",
         y = "log (SD)") + 
    theme_classic()

ggarrange(male, female, ncol = 2, nrow = 1, labels = c("A", "B"))
```

Figure 2.1: Mean-variance relationships (log(Mean) vs log(SD, standard deviation)) across all traits for males (A) and females (B).

We can see that there are strong relationships between the log transformed means and standard deviations for both males (Figure 2.1A) and females (Figure 2.1B), with the Person’s correlation coefficient being 0.949 and 0.95, respectively.

### 3 Meta-analyses

#### 3.1 Population as analysis unit

In this section, we cover the workflow described in Figure 1.1C and D.

##### 3.1.1 Loop through meta-analyses on all traits

This section covers Figure 1.1 C, by conducting a meta-analysis for all traits and producing the meta-analysed effect sizes (lnVR, lnCVR and lnRR).

- The loop combines the functions mentioned above and fills the data matrix with results from our meta analysis.
- Error messages indicate traits that either did not reach convergence, or that did not return meaningful results in the meta-analysis, due to absence of variance. Those traits will be removed in later steps, outlined below.

```
n <- length(unique(data$id))

# Create dataframe to store results
results_alltraits_grouping <- data.frame(tibble(id = 1:n, lnCVR = 0, lnCVR_lower = 0, 
    lnCVR_upper = 0, lnCVR_se = 0, lnCVR_I2 = 0, lnVR = 0, lnVR_lower = 0, lnVR_upper = 0, 
    lnVR_se = 0, lnVR_I2 = 0, lnRR = 0, lnRR_lower = 0, lnRR_upper = 0, lnRR_se = 0, 
    lnRR_I2 = 0, k = 0, trait = 0, male_N = 0, female_N = 0, min_male_N = 0, max_male_N = 0, 
    mean_male_N = 0, min_female_N = 0, max_female_N = 0, mean_female_N = 0))

for (t in 1:n) {
    tryCatch({
        
        results <- data %>% data_subset_parameterid_individual_by_age(t) %>% calculate_population_stats() %>% 
            create_meta_analysis_effect_sizes() %>% mutate(err = seq_len(n()))
        
        # lnCVR, log repsonse-ratio of the coefficient of variance
        cvr <- metafor::rma.mv(yi = effect_size_CVR, V = sample_variance_CVR, random = list(~1 | 
            strain_name, ~1 | production_center, ~1 | err), control = list(optimizer = "optim", 
            optmethod = "Nelder-Mead", maxit = 1000), verbose = F, data = results)
        
        # lnVR, comparison of standard deviations
        cv <- metafor::rma.mv(yi = effect_size_VR, V = sample_variance_VR, random = list(~1 | 
            strain_name, ~1 | production_center, ~1 | err), control = list(optimizer = "optim", 
            optmethod = "Nelder-Mead", maxit = 1000), verbose = F, data = results)
        
        # for means, lnRR
        means <- metafor::rma.mv(yi = effect_size_ROM, V = sample_variance_ROM, random = list(~1 | 
            strain_name, ~1 | production_center, ~1 | err), control = list(optimizer = "optim", 
            optmethod = "Nelder-Mead", maxit = 1000), verbose = F, data = results)
        
        f <- function(x) unlist(x[c("b", "ci.lb", "ci.ub", "se")])
        
        extract_I2 <- function(mod) {
            sigma2 <- mod$sigma2
            w <- mod$vi
            k <- mod$k
            
            sigmaM <- sum(w * (k - 1))/(sum(w)^2 - sum(w^2))
            
            I2_tot <- round(sum(sigma2)/sum(sigma2 + sigmaM), digits = 3) * 100
            return(I2_tot)
        }
        results_alltraits_grouping[t, c(2:17, 19:26)] <- c(f(cvr), lnCVR_I2 = extract_I2(cvr), 
            f(cv), lnVR_I2 = extract_I2(cv), f(means), lnRR_I2 = extract_I2(means), 
            k = means$k, male_N = sum(results$male_n_ind), female_N = sum(results$female_n_ind), 
            min_male_N = min(results$male_n_ind), max_male_N = max(results$male_n_ind), 
            mean_male_N = mean(results$male_n_ind), min_female_N = min(results$female_n_ind), 
            max_female_N = max(results$female_n_ind), mean_female_N = mean(results$female_n_ind))
        results_alltraits_grouping[t, 18] <- unique(results$trait)
    }, error = function(e) {
        cat("ERROR :", t, conditionMessage(e), "\n")
    })
}
```

```
## ERROR : 84 Optimizer (optim) did not achieve convergence (convergence = 10). 
## ERROR : 160 Processing terminated since k <= 1. 
## ERROR : 168 Processing terminated since k <= 1.
```

In the above function, we use ‘tryCatch’ and ‘conditionMessage’ to prevent the loop from aborting when the first error at row 84 is produced. As convergence in the two listed non-converging cases can’t be achieved by sensibly tweaking (other optim etc.), and we only learn about non-convergence in the loop, it is not possible to exclude the traits (N = 2) beforehand.Similarly, there are 8 traits with very low variation, which can not be excluded prior to running the loop.

Any produced “Warnings” indicate cases where variance components are set to zero during likelihood optimization.

##### 3.1.2 Merging datasets & removal of non-converged traits

Here we cover Figure 1.1D, removing non-coverging model results and non-informative traits. Procedure names, grouping variables and trait names (“parameter\_names”) are the merged back together with the results from the metafor analysis above.

```
results_alltraits_grouping2 <- results_alltraits_grouping %>% 
# Join data with results_alltraits_grouping. We filter duplicated id's to get
# only one unique row per id (and there is one id per parameter_name)
left_join(by = "id", data %>% select(id, parameter_group, procedure = procedure_name, 
    procedure_name, parameter_name) %>% filter(!duplicated(id))) %>% 
# Below we add 'procedure' (from the previously loaded 'procedures.csv') as a
# variable; n <- length(unique(results_alltraits_grouping2$parameter_name)))
# should equal 232
left_join(by = "procedure", procedures %>% distinct())

# We exclude 14 parameter names for which metafor models didn't converge ('dp t
# cells', 'mzb (cd21/35 high)'), and of parameters that don't harbour enough
# variation
meta_clean <- results_alltraits_grouping2 %>% filter(!parameter_name %in% c("dp t cells", 
    "mzb (cd21/35 high)", "number of caudal vertebrae", "number of cervical vertebrae", 
    "number of digits", "number of lumbar vertebrae", "number of pelvic vertebrae", 
    "number of ribs left", "number of ribs right", "number of signals", "number of thoracic vertebrae", 
    "total number of acquired events in panel a", "total number of acquired events in panel b", 
    "whole arena permanence"))

# Summary of the cleaned data for 218 traits
sd(meta_clean$k)
range(meta_clean$male_N)
range(meta_clean$female_N)
range(meta_clean$lnCVR_I2)
range(meta_clean$lnVR_I2)
range(meta_clean$lnRR_I2)
range(meta_clean$k)
mean(meta_clean$mean_male_N)
range(meta_clean$min_male_N)
median(meta_clean$mean_male_N)
sd(meta_clean$mean_male_N)
sd(meta_clean$mean_male_N)/sqrt(length(meta_clean$mean_male_N))
mean(meta_clean$mean_female_N)
median(meta_clean$mean_female_N)
range(meta_clean$min_female_N)
sd(meta_clean$mean_female_N)
sd(meta_clean$mean_female_N)/sqrt(length(meta_clean$mean_female_N))
```

A total of 14 traits from the original 232 that had been included are removed because they either did not achieve convergence or are nonsensical for analysis of variance (such as traits that show no variation, see list below).

The traits where models did not converge include: “dp t cells”, “mzb (cd21/35 high)”

The traits where there was not enough variation include: “number of caudal vertebrae”, “number of cervical vertebrae”, “number of digits”, “number of lumbar vertebrae”, “number of pelvic vertebrae”, “number of ribs left”,“number of ribs right”, “number of signals”, “number of thoracic vertebrae”, “total number of acquired events in panel a”,“total number of acquired events in panel b”, “whole arena permanence”.

#### 3.2 Meta-analysis: condensing non-independent traits

This dataset contained a number of highly correlated traits, such as different kinds of cell counts (for example hierarchical parameterization within immunological assays). As those data-points are not independent of each other, we conducted meta analyses on these correlated parameters to collapse the number of levels. This section describes the workflow depicted in Figure 1.1E & F.

##### 3.2.1 Collapsing and merging correlated parameters

Count the number of parameter names (correlated sub-traits) in each parameter group (par\_group\_size). This serves to identify and separate the traits that are correlated from the full dataset that can be processed as is. If the sample size (n) for a given “parameter group” equals 1, the trait is unique and uncorrelated. All instances, where there are 2 or more traits associated with the same parameter group (90 cases), are selected for a “mini-meta analysis”, which removes the issue of correlation. The workflow here relates to Figure 1.1E.

```
# Here we double check numbers of trait parameters in the dataset
meta1 <- meta_clean
# length(unique(meta1$procedure)) #18 length(unique(meta1$GroupingTerm)) #9
# length(unique(meta1$parameter_group)) # 148 levels. To be used as grouping
# factor for meta-meta-analysis / collapsing down based on things that are
# classified identically in 'parameter_group' but have different 'parameter_name'
# length(unique(meta1$parameter_name)) #218 and prepare the dataset for
# second-order meta-analysis
meta1_sub <- meta1 %>% # Add summary of number of parameter names in each parameter group
group_by(parameter_group) %>% mutate(par_group_size = length(unique(parameter_name)), 
    sampleSize = as.numeric(k)) %>% ungroup() %>% # Create subsets with > 1 count (par_group_size > 1)
filter(par_group_size > 1)  # 90 observations
```

##### 3.2.2 Meta-analyses on correlated (sub-)traits, using `robumeta`

Here we pepare the subset of the data (using nest()), and in this first step the model of the meta-analysis effect sizes are calculated. This section specifically undertakes the workflow described in Figure 1.1 F.

```
# Create summary of number of parameter names in each parameter group, and merge
# back together

meta1b <- meta1 %>% group_by(parameter_group) %>% summarize(par_group_size = length(unique(parameter_name, 
    na.rm = TRUE)))

meta1$par_group_size <- meta1b$par_group_size[match(meta1$parameter_group, meta1b$parameter_group)]

# Create subsets with > 1 count (par_group_size > 1)

meta1_sub <- subset(meta1, par_group_size > 1)  # 90 observations   
meta1_sub$sampleSize <- as.numeric(meta1_sub$k)

# Nesting and meta-analyses on correlated traits, using robumeta

n_count <- meta1_sub %>% group_by(parameter_group) %>% mutate(raw_N = sum(sampleSize)) %>% 
    nest() %>% ungroup()

model_count <- n_count %>% mutate(model_lnRR = map(data, ~robu(.x$lnRR ~ 1, data = .x, 
    studynum = .x$id, modelweights = c("CORR"), rho = 0.8, small = TRUE, var.eff.size = (.x$lnRR_se)^2)), 
    model_lnVR = map(data, ~robu(.x$lnVR ~ 1, data = .x, studynum = .x$id, modelweights = c("CORR"), 
        rho = 0.8, small = TRUE, var.eff.size = (.x$lnVR_se)^2)), model_lnCVR = map(data, 
        ~robu(.x$lnCVR ~ 1, data = .x, studynum = .x$id, modelweights = c("CORR"), 
            rho = 0.8, small = TRUE, var.eff.size = (.x$lnCVR_se)^2)))


#### Extracting and save parameter estimates Here we apply an additional function to
#### collect the outcomes of the 'mini-meta-analysis' that has condensed our
#### non-independent traits. Values from our second-order meta-analysis using
#### robu-meta are then extracted

count_fun <- function(mod_sub) {
    return(c(mod_sub$reg_table$b.r, mod_sub$reg_table$CI.L, mod_sub$reg_table$CI.U, 
        mod_sub$reg_table$SE))
}  # estimate, lower ci, upper ci, SE

# Extraction of values created during meta-analyses using robumeta

robusub_RR <- model_count %>% transmute(parameter_group, estimatelnRR = map(model_lnRR, 
    count_fun)) %>% mutate(r = map(estimatelnRR, ~data.frame(t(.)))) %>% unnest(r) %>% 
    select(-estimatelnRR) %>% purrr::set_names(c("parameter_group", "lnRR", "lnRR_lower", 
    "lnRR_upper", "lnRR_se"))

robusub_CVR <- model_count %>% transmute(parameter_group, estimatelnCVR = map(model_lnCVR, 
    count_fun)) %>% mutate(r = map(estimatelnCVR, ~data.frame(t(.)))) %>% unnest(r) %>% 
    select(-estimatelnCVR) %>% purrr::set_names(c("parameter_group", "lnCVR", "lnCVR_lower", 
    "lnCVR_upper", "lnCVR_se"))

robusub_VR <- model_count %>% transmute(parameter_group, estimatelnVR = map(model_lnVR, 
    count_fun)) %>% mutate(r = map(estimatelnVR, ~data.frame(t(.)))) %>% unnest(r) %>% 
    select(-estimatelnVR) %>% purrr::set_names(c("parameter_group", "lnVR", "lnVR_lower", 
    "lnVR_upper", "lnVR_se"))

robu_all <- full_join(robusub_CVR, robusub_VR) %>% full_join(., robusub_RR)

#### Combining data Merge the two data sets (the new [robu_all] and the initial
#### [uncorrelated sub-traits with count = 1])
meta_all <- meta1 %>% filter(par_group_size == 1) %>% as_tibble()
# glimpse(meta_all) glimpse(robu_all)

# Step 1: Columns are matched by name (in our case, 'parameter_group'), and any
# missing columns will be filled with NA
combinedmeta <- bind_rows(robu_all, meta_all)
# glimpse(combinedmeta)

# Steps 2&3: Add information about number of traits in a parameter group,
# procedure, and grouping term
metacombo <- combinedmeta
metacombo$counts <- meta1$par_group_size[match(metacombo$parameter_group, meta1$parameter_group)]
metacombo$procedure2 <- meta1$procedure[match(metacombo$parameter_group, meta1$parameter_group)]
metacombo$GroupingTerm2 <- meta1$GroupingTerm[match(metacombo$parameter_group, meta1$parameter_group)]

# Clean-up, reorder, and rename

metacombo <- metacombo[c("parameter_group", "counts", "procedure2", "GroupingTerm2", 
    "lnCVR", "lnCVR_lower", "lnCVR_upper", "lnCVR_se", "lnVR", "lnVR_lower", "lnVR_upper", 
    "lnVR_se", "lnRR", "lnRR_lower", "lnRR_upper", "lnRR_se")]

names(metacombo)[names(metacombo) == "procedure2"] <- "procedure"
names(metacombo)[names(metacombo) == "GroupingTerm2"] <- "GroupingTerm"
```

#### 3.3 Second-order meta-analysis for functional groups

Here, we describe the workflow depicted in Figure 1.1H. We now use our uncorrelated traits from our first order meta-analysis to conduct a secondary meta-analysis by analysing the 9 functional trait groups. We estimate an overall mean lnCVR, lnVR and lnRR for each functional trait group.

##### 3.3.1 Preparing the data

Nesting, calculating the number of parameters within each grouping term, and running the meta-analyses.

```
metacombo_final <- metacombo %>% group_by(GroupingTerm) %>% nest()

# **Calculate number of parameters per grouping term

metacombo_final <- metacombo_final %>% mutate(para_per_GroupingTerm = map_dbl(data, 
    nrow))

# For all grouping terms
metacombo_final_all <- metacombo %>% nest(data = everything())


# Final random effects meta-analyses within grouping terms, with SE of the
# estimate

overall1 <- metacombo_final %>% 
mutate(model_lnCVR = map(data, ~metafor::rma.uni(yi = .x$lnCVR, sei = (.x$lnCVR_upper - 
    .x$lnCVR_lower)/(2 * 1.96), control = list(optimizer = "optim", optmethod = "Nelder-Mead", 
    maxit = 1000), verbose = F)), model_lnVR = map(data, ~metafor::rma.uni(yi = .x$lnVR, 
    sei = (.x$lnVR_upper - .x$lnVR_lower)/(2 * 1.96), control = list(optimizer = "optim", 
        optmethod = "Nelder-Mead", maxit = 1000), verbose = F)), model_lnRR = map(data, 
    ~metafor::rma.uni(yi = .x$lnRR, sei = (.x$lnRR_upper - .x$lnRR_lower)/(2 * 1.96), 
        control = list(optimizer = "optim", optmethod = "Nelder-Mead", maxit = 1000), 
        verbose = F)))

# Final random effects meta-analyses ACROSS grouping terms, with SE of the
# estimate

overall_all1 <- metacombo_final_all %>% 
mutate(model_lnCVR = map(data, ~metafor::rma.uni(yi = .x$lnCVR, sei = (.x$lnCVR_upper - 
    .x$lnCVR_lower)/(2 * 1.96), control = list(optimizer = "optim", optmethod = "Nelder-Mead", 
    maxit = 1000), verbose = F)), model_lnVR = map(data, ~metafor::rma.uni(yi = .x$lnVR, 
    sei = (.x$lnVR_upper - .x$lnVR_lower)/(2 * 1.96), control = list(optimizer = "optim", 
        optmethod = "Nelder-Mead", maxit = 1000), verbose = F)), model_lnRR = map(data, 
    ~metafor::rma.uni(yi = .x$lnRR, sei = (.x$lnRR_upper - .x$lnRR_lower)/(2 * 1.96), 
        control = list(optimizer = "optim", optmethod = "Nelder-Mead", maxit = 1000), 
        verbose = F)))
```

##### 3.3.2 Re-structuring the data for each grouping term

Here we delete unused variables, and select the respective effect sizes. Please note - the referencing of the cells does NOT depend on previous ordering of the data. This would only be affected if the output structure from metafor::rma.uni changes.

```
# Function will take the averall results and extract the relevant trait of
# interest
extract_trait <- function(data, trait) {
    tmp <- data %>% filter(., GroupingTerm == trait) %>% mutate(lnCVR = .[[4]][[1]]$b, 
        lnCVR_lower = .[[4]][[1]]$ci.lb, lnCVR_upper = .[[4]][[1]]$ci.ub, lnCVR_se = .[[4]][[1]]$se, 
        lnCVR_I2 = .[[4]][[1]]$I2, lnVR = .[[5]][[1]]$b, lnVR_lower = .[[5]][[1]]$ci.lb, 
        lnVR_upper = .[[5]][[1]]$ci.ub, lnVR_se = .[[5]][[1]]$se, lnVR_I2 = .[[5]][[1]]$I2, 
        lnRR = .[[6]][[1]]$b, lnRR_lower = .[[6]][[1]]$ci.lb, lnRR_upper = .[[6]][[1]]$ci.ub, 
        lnRR_se = .[[6]][[1]]$se, lnRR_I2 = .[[6]][[1]]$I2) %>% select(., GroupingTerm, 
        lnCVR:lnRR_I2)
    
    return(tmp)
}

All <- overall_all1 %>% mutate(lnCVR = .[[2]][[1]]$b, lnCVR_lower = .[[2]][[1]]$ci.lb, 
    lnCVR_upper = .[[2]][[1]]$ci.ub, lnCVR_se = .[[2]][[1]]$se, lnCVR_I2 = .[[2]][[1]]$I2, 
    lnVR = .[[3]][[1]]$b, lnVR_lower = .[[3]][[1]]$ci.lb, lnVR_upper = .[[3]][[1]]$ci.ub, 
    lnVR_se = .[[3]][[1]]$se, lnVR_I2 = .[[3]][[1]]$I2, lnRR = .[[4]][[1]]$b, lnRR_lower = .[[4]][[1]]$ci.lb, 
    lnRR_upper = .[[4]][[1]]$ci.ub, lnRR_se = .[[4]][[1]]$se, lnRR_I2 = .[[4]][[1]]$I2, 
    ) %>% select(., lnCVR:lnRR_I2)

All <- All %>% mutate(GroupingTerm = "All")

overall2 <- bind_rows(extract_trait(overall1, "Behaviour"), extract_trait(overall1, 
    "Morphology"), extract_trait(overall1, "Metabolism"), extract_trait(overall1, 
    "Physiology"), extract_trait(overall1, "Immunology"), extract_trait(overall1, 
    "Hematology"), extract_trait(overall1, "Heart"), extract_trait(overall1, "Hearing"), 
    extract_trait(overall1, "Eye"), All)
```

#### 3.4 Second-order meta-analysis of heterogenity

To see how much heterogeneity there is for each trait across different populations on average (for different functional groups and overall), we also conducted secondary meta-analyses by combining the total heterogeneities. More specifically, we combined the logarithm of the sum of the three variance components (Strain, Location and Unit; see above) with 1/2(N[effect size] - 1) as the sampling variance via the random effect model. That is, we conducted 3 sets of 10 secondary meta-analyses as with those of mean effect sizes for each trait, i.e. meta-analyzing log(total heterogeneities) for 9 functional groups and all the groups combined (Figure 1.1I). The analysis for heterogeneity follows the workflow of the above steps for the different meta-analyses and specifically Figure 1.1I & J. However, in the initial meta-analysis we extract sigma^2 and errors for mouse strains and centers (Institutions). As above a Loop is run and parameters extracted from metafor (sigma2’s, s.nlevels)

```
### Preparing heterogeneity

# Create dataframe to store results
results.allhetero.grouping <- as.data.frame(cbind(c(1:n), matrix(rep(0, n * 30), 
    ncol = 30)))
names(results.allhetero.grouping) <- c("id", "sigma2_strain.CVR", "sigma2_center.CVR", 
    "sigma2_error.CVR", "s.nlevels.strain.CVR", "s.nlevels.center.CVR", "s.nlevels.error.CVR", 
    "sigma2_strain.VR", "sigma2_center.VR", "sigma2_error.VR", "s.nlevels.strain.VR", 
    "s.nlevels.center.VR", "s.nlevels.error.VR", "sigma2_strain.RR", "sigma2_center.RR", 
    "sigma2_error.RR", "s.nlevels.strain.RR", "s.nlevels.center.RR", "s.nlevels.error.RR", 
    "lnCVR", "lnCVR_lower", "lnCVR_upper", "lnCVR_se", "lnVR", "lnVR_lower", "lnVR_upper", 
    "lnVR_se", "lnRR", "lnRR_lower", "lnRR_upper", "lnRR_se")

# Loop

for (t in 1:n) {
    tryCatch({
        results <- data %>% data_subset_parameterid_individual_by_age(t) %>% calculate_population_stats() %>% 
            create_meta_analysis_effect_sizes() %>% mutate(err = seq_len(n()))
        
        
        # lnCVR, logaritm of the ratio of male and female coefficients of variance
        
        cvr. <- metafor::rma.mv(yi = effect_size_CVR, V = sample_variance_CVR, random = list(~1 | 
            strain_name, ~1 | production_center, ~1 | err), control = list(optimizer = "optim", 
            optmethod = "Nelder-Mead", maxit = 1000), data = results)
        
        results.allhetero.grouping[t, 2] <- cvr.$sigma2[1]
        results.allhetero.grouping[t, 3] <- cvr.$sigma2[2]
        results.allhetero.grouping[t, 4] <- cvr.$sigma2[3]
        results.allhetero.grouping[t, 5] <- cvr.$s.nlevels[1]
        results.allhetero.grouping[t, 6] <- cvr.$s.nlevels[2]
        results.allhetero.grouping[t, 7] <- cvr.$s.nlevels[3]
        results.allhetero.grouping[t, 20] <- cvr.$b
        results.allhetero.grouping[t, 21] <- cvr.$ci.lb
        results.allhetero.grouping[t, 22] <- cvr.$ci.ub
        results.allhetero.grouping[t, 23] <- cvr.$se
        
        # lnVR, male to female variability ratio (logarithm of male and female standard
        # deviations)
        
        vr. <- metafor::rma.mv(yi = effect_size_VR, V = sample_variance_VR, random = list(~1 | 
            strain_name, ~1 | production_center, ~1 | err), control = list(optimizer = "optim", 
            optmethod = "Nelder-Mead", maxit = 1000), data = results)
        
        results.allhetero.grouping[t, 8] <- vr.$sigma2[1]
        results.allhetero.grouping[t, 9] <- vr.$sigma2[2]
        results.allhetero.grouping[t, 10] <- vr.$sigma2[3]
        results.allhetero.grouping[t, 11] <- vr.$s.nlevels[1]
        results.allhetero.grouping[t, 12] <- vr.$s.nlevels[2]
        results.allhetero.grouping[t, 13] <- vr.$s.nlevels[3]
        results.allhetero.grouping[t, 24] <- vr.$b
        results.allhetero.grouping[t, 25] <- vr.$ci.lb
        results.allhetero.grouping[t, 26] <- vr.$ci.ub
        results.allhetero.grouping[t, 27] <- vr.$se
        
        # lnRR, response ratio (logarithm of male and female means)
        
        rr. <- metafor::rma.mv(yi = effect_size_ROM, V = sample_variance_ROM, random = list(~1 | 
            strain_name, ~1 | production_center, ~1 | err), control = list(optimizer = "optim", 
            optmethod = "Nelder-Mead", maxit = 1000), data = results)
        
        results.allhetero.grouping[t, 14] <- rr.$sigma2[1]
        results.allhetero.grouping[t, 15] <- rr.$sigma2[2]
        results.allhetero.grouping[t, 16] <- rr.$sigma2[3]
        results.allhetero.grouping[t, 17] <- rr.$s.nlevels[1]
        results.allhetero.grouping[t, 18] <- rr.$s.nlevels[2]
        results.allhetero.grouping[t, 19] <- rr.$s.nlevels[3]
        results.allhetero.grouping[t, 28] <- rr.$b
        results.allhetero.grouping[t, 29] <- rr.$ci.lb
        results.allhetero.grouping[t, 30] <- rr.$ci.ub
        results.allhetero.grouping[t, 31] <- rr.$se
    }, error = function(e) {
        cat("ERROR :", conditionMessage(e), "\n")
    })
}
```

```
## ERROR : Optimizer (optim) did not achieve convergence (convergence = 10). 
## ERROR : Processing terminated since k <= 1. 
## ERROR : Processing terminated since k <= 1.
```

Here we exclude traits without variation between mouse strains and merge data sets containing metafor results with procedure etc. names. Dealing with the correlated parameters, and running the second -order meta-analysis for heterogeneity data (Figure 1.1I)

```
              results.allhetero.grouping2 <- results.allhetero.grouping[results.allhetero.grouping$s.nlevels.strain.VR != 0, ]
# nrow(results.allhetero.grouping) #232  

# Merging dataset with correct procedure names
# procedures <- read.csv(here("export", "procedures.csv"))

results.allhetero.grouping2$parameter_group <- data$parameter_group[match(results.allhetero.grouping2$id, data$id)]
      results.allhetero.grouping2$procedure <- data$procedure_name[match(results.allhetero.grouping2$id, data$id)]

   results.allhetero.grouping2$GroupingTerm <- procedures$GroupingTerm[match(results.allhetero.grouping2$procedure, procedures$procedure)]
 results.allhetero.grouping2$parameter_name <- data$parameter_name[match(results.allhetero.grouping2$id, data$id)]

#We deal with correlated parameters following the procedures described above (extraction of effects sizes).

metahetero1 <- results.allhetero.grouping2
# length(unique(metahetero1$procedure)) #19
# length(unique(metahetero1$GroupingTerm)) #9 
# length(unique(metahetero1$parameter_group)) #152
# length(unique(metahetero1$parameter_name)) #223

# Count of number of parameter names (correlated sub-traits) in each parameter group (par_group_size)

metahetero1b <- metahetero1                    %>%
                group_by(parameter_group)      %>%
                mutate(par_group_size = n_distinct(parameter_name))

metahetero1$par_group_size <- metahetero1b$par_group_size[match(metahetero1$parameter_group, metahetero1b$parameter_group)]

# Create subsets with > 1 count (par_group_size > 1)

metahetero1_sub <- subset(metahetero1, par_group_size > 1) # 92 observations

# Nest data

n_count. <- metahetero1_sub %>%
  group_by(parameter_group) %>%
  nest()

# meta-analysis preparation

model_count. <- n_count. %>%
  mutate(
    model_lnRR = map(data, ~ robu(.x$lnRR ~ 1,
      data = .x, studynum = .x$id, modelweights = c("CORR"), rho = 0.8,
      small = TRUE, var.eff.size = (.x$lnRR_se)^2
    )),
    model_lnVR = map(data, ~ robu(.x$lnVR ~ 1,
      data = .x, studynum = .x$id, modelweights = c("CORR"), rho = 0.8,
      small = TRUE, var.eff.size = (.x$lnVR_se)^2
    )),
    model_lnCVR = map(data, ~ robu(.x$lnCVR ~ 1,
      data = .x, studynum = .x$id, modelweights = c("CORR"), rho = 0.8,
      small = TRUE, var.eff.size = (.x$lnCVR_se)^2
    ))
  )


# Robumeta object details:  str(model_count.$model_lnCVR[[1]])

# Perform meta-analyses on correlated sub-traits, using robumeta
# Extract and save parameter estimates

count_fun. <- function(mod_sub) {
  return(c(as.numeric(mod_sub$mod_info$term1), mod_sub$N))
}

robusub_RR. <- model_count. %>%
  transmute(estimatelnRR = map(model_lnRR, count_fun.)) %>%    
  mutate(r = map(estimatelnRR, ~ data.frame(t(.)))) %>%
  unnest(r) %>%
  select(-estimatelnRR) %>%
  purrr::set_names(c("parameter_group", "var.RR", "N.RR"))

robusub_CVR. <- model_count. %>%
  transmute(estimatelnCVR = map(model_lnCVR, count_fun.)) %>%
  mutate(r = map(estimatelnCVR, ~ data.frame(t(.)))) %>%
  unnest(r) %>%
  select(-estimatelnCVR) %>%
  purrr::set_names(c("parameter_group", "var.CVR", "N.CVR"))

robusub_VR. <- model_count. %>%
  transmute(estimatelnVR = map(model_lnVR, count_fun.)) %>%
  mutate(r = map(estimatelnVR, ~ data.frame(t(.)))) %>%
  unnest(r) %>%
  select(-estimatelnVR) %>%
  purrr::set_names(c("parameter_group", "var.VR", "N.VR"))

robu_all. <- full_join(robusub_CVR., robusub_VR.) %>% full_join(., robusub_RR.)


#Merge the two data sets (the new [robu_all.] and the initial [uncorrelated sub-traits with count = 1])

#In this step, we   1) merge the N from robumeta and the  N from metafor (s.nlevels.error) together into the same columns (N.RR, N.VR, N.CVR)

# 2) calculate the total variance for metafor models as the sum of random effect variances and the residual error, then add in the same columns together with the residual variances from robumeta 
  
  
metahetero_all <- metahetero1 %>%
  filter(par_group_size == 1) %>%
  as_tibble()
metahetero_all$N.RR <- metahetero_all$s.nlevels.error.RR
metahetero_all$N.CVR <- metahetero_all$s.nlevels.error.CVR
metahetero_all$N.VR <- metahetero_all$s.nlevels.error.VR
metahetero_all$var.RR <- log(sqrt(metahetero_all$sigma2_strain.RR + metahetero_all$sigma2_center.RR + metahetero_all$sigma2_error.RR))
metahetero_all$var.VR <- log(sqrt(metahetero_all$sigma2_strain.VR + metahetero_all$sigma2_center.VR + metahetero_all$sigma2_error.VR))
metahetero_all$var.CVR <- log(sqrt(metahetero_all$sigma2_strain.CVR + metahetero_all$sigma2_center.CVR + metahetero_all$sigma2_error.CVR))
# str(metahetero_all)
# str(robu_all.)

metahetero_all <- metahetero_all %>% mutate(
  var.RR = if_else(var.RR == -Inf, -7, var.RR),   # numbers chosen as limits; based on the values in the dataset
  var.VR = if_else(var.VR == -Inf, -5, var.VR),
  var.CVR = if_else(var.CVR == -Inf, -6, var.CVR)
)

# Combine data
# Step1

combinedmetahetero <- bind_rows(robu_all., metahetero_all)
# glimpse(combinedmetahetero)

# Steps 2&3

              metacombohetero <- combinedmetahetero
       metacombohetero$counts <- metahetero1$par_group_size[match(metacombohetero$parameter_group, metahetero1$parameter_group)]
   metacombohetero$procedure2 <- metahetero1$procedure[match(metacombohetero$parameter_group, metahetero1$parameter_group)]
metacombohetero$GroupingTerm2 <- metahetero1$GroupingTerm[match(metacombohetero$parameter_group, metahetero1$parameter_group)]

# **Clean-up and rename

metacombohetero <- metacombohetero %>% select(parameter_group, var.CVR, N.CVR, var.VR, N.VR, var.RR, N.RR, counts, procedure = procedure2, GroupingTerm = GroupingTerm2)  

# Invidual table for heterogeneity across all traits
 kable(metacombohetero,, digits = 3) %>%
  kable_styling() %>%
  scroll_box(width = "100%", height = "200px")
```

| parameter\_group | var.CVR | N.CVR | var.VR | N.VR | var.RR | N.RR | counts | procedure | GroupingTerm |
| --- | --- | --- | --- | --- | --- | --- | --- | --- | --- |
| pre-pulse inhibition | 0.004 | 5 | 0.000 | 5 | 0.000 | 5 | 5 | Acoustic Startle and Pre-pulse Inhibition (PPI) | Behaviour |
| B cells | 0.000 | 4 | 0.000 | 4 | 0.002 | 4 | 4 | Immunophenotyping | Immunology |
| cd4 nkt | -0.014 | 6 | -0.012 | 6 | 0.018 | 6 | 6 | Immunophenotyping | Immunology |
| cd4 t | 0.005 | 7 | 0.003 | 7 | 0.001 | 7 | 7 | Immunophenotyping | Immunology |
| cd8 nkt | -0.008 | 6 | 0.003 | 6 | -0.003 | 6 | 6 | Immunophenotyping | Immunology |
| cd8 t | 0.004 | 7 | 0.001 | 7 | 0.000 | 7 | 7 | Immunophenotyping | Immunology |
| cdcs | -0.002 | 2 | -0.009 | 2 | -0.002 | 2 | 2 | Immunophenotyping | Immunology |
| dn nkt | -0.001 | 6 | 0.001 | 6 | 0.006 | 6 | 6 | Immunophenotyping | Immunology |
| dn t | 0.005 | 7 | 0.000 | 7 | 0.000 | 7 | 7 | Immunophenotyping | Immunology |
| eosinophils | 0.000 | 3 | 0.013 | 3 | 0.005 | 3 | 3 | Hematology | Hematology |
| follicular b cells | -0.002 | 2 | -0.007 | 2 | 0.000 | 2 | 2 | Immunophenotyping | Immunology |
| luc | 0.000 | 2 | 0.021 | 2 | 0.027 | 2 | 2 | Hematology | Hematology |
| lymphocytes | 0.059 | 2 | 0.008 | 2 | 0.013 | 2 | 2 | Hematology | Hematology |
| monocytes | 0.002 | 3 | 0.006 | 3 | 0.009 | 3 | 3 | Hematology | Hematology |
| neutrophils | -0.010 | 3 | 0.030 | 3 | 0.018 | 3 | 3 | Hematology | Hematology |
| nk cells | 0.000 | 6 | 0.002 | 6 | 0.002 | 6 | 6 | Immunophenotyping | Immunology |
| nkt cells | 0.000 | 4 | -0.004 | 4 | -0.001 | 4 | 4 | Immunophenotyping | Immunology |
| percentage of live gated events | -0.007 | 2 | -0.004 | 2 | -0.001 | 2 | 2 | Immunophenotyping | Immunology |
| response amplitude | 0.003 | 10 | 0.004 | 10 | 0.013 | 10 | 10 | Acoustic Startle and Pre-pulse Inhibition (PPI) | Behaviour |
| t cells | 0.001 | 3 | 0.011 | 3 | 0.002 | 3 | 3 | Immunophenotyping | Immunology |
| number events | 0.001 | 2 | 0.000 | 2 | -0.001 | 2 | 2 | Immunophenotyping | Immunology |
| 12khz-evoked abr threshold | -2.660 | 14 | -2.144 | 14 | -3.788 | 14 | 1 | Auditory Brain Stem Response | Hearing |
| 18khz-evoked abr threshold | -2.711 | 14 | -2.193 | 14 | -3.514 | 14 | 1 | Auditory Brain Stem Response | Hearing |
| 24khz-evoked abr threshold | -2.467 | 14 | -1.182 | 14 | -3.555 | 14 | 1 | Auditory Brain Stem Response | Hearing |
| 30khz-evoked abr threshold | -2.419 | 14 | -2.456 | 14 | -3.401 | 14 | 1 | Auditory Brain Stem Response | Hearing |
| 6khz-evoked abr threshold | -6.000 | 14 | -4.570 | 14 | -4.675 | 14 | 1 | Auditory Brain Stem Response | Hearing |
| alanine aminotransferase | -1.677 | 16 | -1.116 | 16 | -2.143 | 16 | 1 | Clinical Chemistry | Physiology |
| albumin | -2.221 | 16 | -2.302 | 16 | -3.575 | 16 | 1 | Clinical Chemistry | Physiology |
| alkaline phosphatase | -2.480 | 16 | -2.125 | 16 | -2.553 | 16 | 1 | Clinical Chemistry | Physiology |
| alpha-amylase | -2.316 | 9 | -1.846 | 9 | -3.002 | 9 | 1 | Clinical Chemistry | Physiology |
| area under glucose response curve | -2.146 | 16 | -2.088 | 16 | -2.192 | 16 | 1 | Intraperitoneal glucose tolerance test (IPGTT) | Metabolism |
| aspartate aminotransferase | -1.487 | 16 | -1.111 | 16 | -2.123 | 16 | 1 | Clinical Chemistry | Physiology |
| basophil cell count | -2.440 | 8 | -1.278 | 8 | -1.527 | 8 | 1 | Hematology | Hematology |
| basophil differential count | -2.659 | 8 | -1.253 | 8 | -2.318 | 8 | 1 | Hematology | Hematology |
| bmc/body weight | -1.871 | 17 | -2.057 | 17 | -2.580 | 17 | 1 | Body Composition (DEXA lean/fat) | Morphology |
| body length | -2.749 | 16 | -2.780 | 16 | -4.854 | 16 | 1 | Body Composition (DEXA lean/fat) | Morphology |
| body temp | -6.000 | 3 | -5.000 | 3 | -6.576 | 3 | 1 | Echo | Heart |
| body weight | -2.210 | 18 | -2.251 | 18 | -3.638 | 18 | 1 | Body Weight | Morphology |
| body weight after experiment | -2.837 | 9 | -2.949 | 9 | -3.828 | 9 | 1 | Indirect Calorimetry | Metabolism |
| body weight before experiment | -2.597 | 9 | -2.688 | 9 | -3.677 | 9 | 1 | Indirect Calorimetry | Metabolism |
| bone area | -2.037 | 17 | -1.964 | 17 | -3.014 | 17 | 1 | Body Composition (DEXA lean/fat) | Morphology |
| bone mineral content (excluding skull) | -1.799 | 17 | -1.777 | 17 | -2.565 | 17 | 1 | Body Composition (DEXA lean/fat) | Morphology |
| bone mineral density (excluding skull) | -1.324 | 17 | -1.230 | 17 | -3.506 | 17 | 1 | Body Composition (DEXA lean/fat) | Morphology |
| calcium | -2.447 | 16 | -2.448 | 16 | -5.041 | 16 | 1 | Clinical Chemistry | Physiology |
| cardiac output | -2.671 | 5 | -2.847 | 5 | -3.575 | 5 | 1 | Echo | Heart |
| center average speed | -2.564 | 12 | -2.981 | 12 | -2.753 | 12 | 1 | Open Field | Behaviour |
| center distance travelled | -2.777 | 12 | -1.859 | 12 | -1.859 | 12 | 1 | Open Field | Behaviour |
| center permanence time | -2.852 | 13 | -2.078 | 13 | -1.972 | 13 | 1 | Open Field | Behaviour |
| center resting time | -2.418 | 10 | -1.752 | 10 | -1.576 | 10 | 1 | Open Field | Behaviour |
| chloride | -1.617 | 10 | -1.565 | 10 | -5.111 | 10 | 1 | Clinical Chemistry | Physiology |
| click-evoked abr threshold | -2.317 | 11 | -1.793 | 11 | -2.889 | 11 | 1 | Auditory Brain Stem Response | Hearing |
| creatine kinase | -2.138 | 9 | -1.168 | 9 | -1.340 | 9 | 1 | Clinical Chemistry | Physiology |
| creatinine | -2.358 | 16 | -0.614 | 16 | -2.654 | 16 | 1 | Clinical Chemistry | Physiology |
| cv | -2.160 | 9 | -1.615 | 9 | -2.083 | 9 | 1 | Electrocardiogram (ECG) | Heart |
| distance travelled - total | -2.312 | 13 | -2.212 | 13 | -2.242 | 13 | 1 | Open Field | Behaviour |
| ejection fraction | -2.477 | 6 | -2.588 | 6 | -4.039 | 6 | 1 | Echo | Heart |
| end-diastolic diameter | -6.000 | 3 | -3.219 | 3 | -4.076 | 3 | 1 | Echo | Heart |
| end-systolic diameter | -6.000 | 3 | -5.000 | 3 | -3.875 | 3 | 1 | Echo | Heart |
| fasted blood glucose concentration | -1.978 | 16 | -2.041 | 16 | -2.915 | 16 | 1 | Intraperitoneal glucose tolerance test (IPGTT) | Metabolism |
| fat mass | -2.007 | 17 | -1.755 | 17 | -2.267 | 17 | 1 | Body Composition (DEXA lean/fat) | Morphology |
| fat/body weight | -1.918 | 17 | -1.806 | 17 | -2.283 | 17 | 1 | Body Composition (DEXA lean/fat) | Morphology |
| forelimb and hindlimb grip strength measurement mean | -2.365 | 16 | -2.197 | 16 | -3.135 | 16 | 1 | Grip Strength | Morphology |
| forelimb grip strength measurement mean | -2.601 | 16 | -2.573 | 16 | -3.234 | 16 | 1 | Grip Strength | Morphology |
| fractional shortening | -2.490 | 7 | -2.520 | 7 | -4.428 | 7 | 1 | Echo | Heart |
| free fatty acids | -2.245 | 5 | -2.204 | 5 | -4.054 | 5 | 1 | Clinical Chemistry | Physiology |
| fructosamine | -2.828 | 6 | -2.795 | 6 | -3.336 | 6 | 1 | Clinical Chemistry | Physiology |
| glucose | -2.546 | 16 | -1.963 | 16 | -2.641 | 16 | 1 | Clinical Chemistry | Physiology |
| hdl-cholesterol | -2.294 | 15 | -2.035 | 15 | -2.612 | 15 | 1 | Clinical Chemistry | Physiology |
| heart weight | -1.978 | 15 | -1.661 | 15 | -3.140 | 15 | 1 | Heart Weight | Morphology |
| heart weight normalised against body weight | -2.199 | 14 | -1.782 | 14 | -3.132 | 14 | 1 | Heart Weight | Morphology |
| hematocrit | -1.578 | 17 | -1.597 | 17 | -3.881 | 17 | 1 | Hematology | Hematology |
| hemoglobin | -2.449 | 17 | -2.389 | 17 | -3.875 | 17 | 1 | Hematology | Hematology |
| hr | -2.090 | 12 | -1.834 | 12 | -2.476 | 12 | 1 | Electrocardiogram (ECG) | Heart |
| hrv | -3.097 | 7 | -1.989 | 7 | -2.003 | 7 | 1 | Electrocardiogram (ECG) | Heart |
| initial response to glucose challenge | -2.842 | 16 | -3.311 | 16 | -2.845 | 16 | 1 | Intraperitoneal glucose tolerance test (IPGTT) | Metabolism |
| insulin | -1.138 | 8 | -0.657 | 8 | -1.041 | 8 | 1 | Insulin Blood Level | Metabolism |
| iron | -1.934 | 11 | -1.739 | 11 | -2.948 | 11 | 1 | Clinical Chemistry | Physiology |
| lactate dehydrogenase | -2.606 | 4 | -1.765 | 4 | -1.989 | 4 | 1 | Clinical Chemistry | Physiology |
| latency to center entry | -2.294 | 10 | -1.555 | 10 | -1.435 | 10 | 1 | Open Field | Behaviour |
| ldl-cholesterol | -1.447 | 7 | -1.147 | 7 | -0.946 | 7 | 1 | Clinical Chemistry | Physiology |
| lean mass | -2.219 | 17 | -2.145 | 17 | -3.418 | 17 | 1 | Body Composition (DEXA lean/fat) | Morphology |
| lean/body weight | -1.719 | 17 | -1.798 | 17 | -3.767 | 17 | 1 | Body Composition (DEXA lean/fat) | Morphology |
| left anterior chamber depth | -1.863 | 2 | -1.852 | 2 | -7.000 | 2 | 1 | Eye Morphology | Eye |
| left corneal thickness | -6.000 | 2 | -5.000 | 2 | -7.000 | 2 | 1 | Eye Morphology | Eye |
| left inner nuclear layer | -6.000 | 2 | -5.000 | 2 | -7.000 | 2 | 1 | Eye Morphology | Eye |
| left outer nuclear layer | -6.000 | 2 | -5.000 | 2 | -7.000 | 2 | 1 | Eye Morphology | Eye |
| left posterior chamber depth | -6.000 | 2 | -5.000 | 2 | -4.923 | 2 | 1 | Eye Morphology | Eye |
| left total retinal thickness | -1.607 | 3 | -1.641 | 3 | -5.300 | 3 | 1 | Eye Morphology | Eye |
| locomotor activity | -2.301 | 13 | -2.882 | 13 | -2.361 | 13 | 1 | Combined SHIRPA and Dysmorphology | Behaviour |
| lvawd | -6.000 | 5 | -5.000 | 5 | -4.648 | 5 | 1 | Echo | Heart |
| lvaws | -1.627 | 4 | -1.885 | 4 | -3.427 | 4 | 1 | Echo | Heart |
| lvidd | -2.763 | 7 | -2.609 | 7 | -4.135 | 7 | 1 | Echo | Heart |
| lvids | -2.137 | 7 | -2.075 | 7 | -4.024 | 7 | 1 | Echo | Heart |
| lvpwd | -2.578 | 7 | -2.307 | 7 | -4.028 | 7 | 1 | Echo | Heart |
| lvpws | -2.718 | 6 | -2.563 | 6 | -3.982 | 6 | 1 | Echo | Heart |
| magnesium | -6.000 | 6 | -3.327 | 6 | -2.767 | 6 | 1 | Urinalysis | Physiology |
| mean cell hemoglobin concentration | -1.404 | 17 | -1.334 | 17 | -5.254 | 17 | 1 | Hematology | Hematology |
| mean cell volume | -1.702 | 17 | -1.737 | 17 | -5.137 | 17 | 1 | Hematology | Hematology |
| mean corpuscular hemoglobin | -2.334 | 17 | -2.325 | 17 | -5.730 | 17 | 1 | Hematology | Hematology |
| mean platelet volume | -2.727 | 13 | -2.650 | 13 | -4.354 | 13 | 1 | Hematology | Hematology |
| mean r amplitude | -6.000 | 8 | -2.751 | 8 | -2.519 | 8 | 1 | Electrocardiogram (ECG) | Heart |
| mean sr amplitude | -4.105 | 7 | -3.587 | 7 | -3.330 | 7 | 1 | Electrocardiogram (ECG) | Heart |
| mzb (cd21/35 high) | -2.182 | 4 | -1.822 | 4 | -7.000 | 4 | 1 | Immunophenotyping | Immunology |
| number of caudal vertebrae | -1.333 | 7 | -1.331 | 7 | -16.871 | 11 | 1 | X-ray | Morphology |
| number of center entries | -2.587 | 10 | -2.550 | 10 | -1.942 | 10 | 1 | Open Field | Behaviour |
| number of digits | -1.530 | 3 | -1.530 | 3 | -16.871 | 15 | 1 | X-ray | Morphology |
| number of lumbar vertebrae | -0.951 | 3 | -0.951 | 3 | -16.871 | 14 | 1 | X-ray | Morphology |
| number of pelvic vertebrae | -1.655 | 3 | -1.655 | 3 | -16.871 | 14 | 1 | X-ray | Morphology |
| number of rears - total | -2.017 | 8 | -1.274 | 8 | -1.892 | 8 | 1 | Open Field | Behaviour |
| number of ribs left | -1.234 | 3 | -1.233 | 3 | -16.871 | 15 | 1 | X-ray | Morphology |
| number of ribs right | -2.400 | 2 | -2.402 | 2 | -16.871 | 15 | 1 | X-ray | Morphology |
| number of signals | -3.327 | 9 | -3.870 | 9 | -3.407 | 11 | 1 | Electrocardiogram (ECG) | Heart |
| others | -3.208 | 6 | -2.931 | 6 | -5.111 | 6 | 1 | Immunophenotyping | Immunology |
| pdcs | -1.637 | 5 | -0.679 | 5 | -1.979 | 5 | 1 | Immunophenotyping | Immunology |
| percentage center time | -2.726 | 13 | -2.123 | 13 | -1.931 | 13 | 1 | Open Field | Behaviour |
| periphery average speed | -2.423 | 12 | -2.283 | 12 | -2.614 | 12 | 1 | Open Field | Behaviour |
| periphery distance travelled | -2.477 | 12 | -2.624 | 12 | -2.273 | 12 | 1 | Open Field | Behaviour |
| periphery permanence time | -1.889 | 13 | -2.136 | 13 | -3.343 | 13 | 1 | Open Field | Behaviour |
| periphery resting time | -2.418 | 10 | -2.895 | 10 | -2.544 | 10 | 1 | Open Field | Behaviour |
| phosphorus | -3.180 | 16 | -2.391 | 16 | -2.927 | 16 | 1 | Clinical Chemistry | Physiology |
| platelet count | -2.517 | 17 | -2.448 | 17 | -3.239 | 17 | 1 | Hematology | Hematology |
| pnn5(6>ms) | -6.000 | 6 | -1.355 | 6 | -1.160 | 6 | 1 | Electrocardiogram (ECG) | Heart |
| potassium | -1.788 | 10 | -1.685 | 10 | -3.393 | 10 | 1 | Clinical Chemistry | Physiology |
| pq | -2.365 | 7 | -2.799 | 7 | -3.489 | 7 | 1 | Electrocardiogram (ECG) | Heart |
| pr | -2.816 | 11 | -2.779 | 11 | -3.976 | 11 | 1 | Electrocardiogram (ECG) | Heart |
| qrs | -3.265 | 11 | -3.212 | 11 | -4.590 | 11 | 1 | Electrocardiogram (ECG) | Heart |
| qtc | -3.176 | 10 | -2.941 | 10 | -4.831 | 10 | 1 | Electrocardiogram (ECG) | Heart |
| qtc dispersion | -2.997 | 7 | -2.265 | 7 | -3.126 | 7 | 1 | Electrocardiogram (ECG) | Heart |
| red blood cell count | -2.269 | 17 | -2.315 | 17 | -3.795 | 17 | 1 | Hematology | Hematology |
| red blood cell distribution width | -1.512 | 13 | -1.425 | 13 | -3.810 | 13 | 1 | Hematology | Hematology |
| respiration rate | -2.972 | 4 | -2.405 | 4 | -3.468 | 4 | 1 | Echo | Heart |
| respiratory exchange ratio | -2.339 | 9 | -2.359 | 9 | -4.825 | 9 | 1 | Indirect Calorimetry | Metabolism |
| right anterior chamber depth | -0.465 | 2 | -0.469 | 2 | -7.000 | 2 | 1 | Eye Morphology | Eye |
| right corneal thickness | -2.191 | 2 | -2.409 | 2 | -4.423 | 2 | 1 | Eye Morphology | Eye |
| right inner nuclear layer | -1.035 | 2 | -0.935 | 2 | -3.449 | 2 | 1 | Eye Morphology | Eye |
| right outer nuclear layer | -6.000 | 2 | -5.000 | 2 | -7.000 | 2 | 1 | Eye Morphology | Eye |
| right posterior chamber depth | -6.000 | 2 | -5.000 | 2 | -4.092 | 2 | 1 | Eye Morphology | Eye |
| right total retinal thickness | -0.948 | 3 | -0.974 | 3 | -4.789 | 3 | 1 | Eye Morphology | Eye |
| rmssd | -1.249 | 7 | -0.802 | 7 | -1.936 | 7 | 1 | Electrocardiogram (ECG) | Heart |
| rp macrophage (cd19- cd11c-) | -1.316 | 6 | -1.285 | 6 | -2.224 | 6 | 1 | Immunophenotyping | Immunology |
| rr | -2.033 | 11 | -1.988 | 11 | -4.619 | 11 | 1 | Electrocardiogram (ECG) | Heart |
| sodium | -1.627 | 10 | -1.544 | 10 | -5.603 | 10 | 1 | Clinical Chemistry | Physiology |
| spleen weight | -1.306 | 10 | -1.020 | 10 | -2.743 | 10 | 1 | Immunophenotyping | Immunology |
| st | -2.830 | 11 | -2.548 | 11 | -4.127 | 11 | 1 | Electrocardiogram (ECG) | Heart |
| stroke volume | -2.179 | 5 | -2.091 | 5 | -4.875 | 5 | 1 | Echo | Heart |
| tibia length | -0.830 | 12 | -0.841 | 12 | -5.345 | 12 | 1 | Heart Weight | Morphology |
| total bilirubin | -2.143 | 16 | -1.809 | 16 | -2.661 | 16 | 1 | Clinical Chemistry | Physiology |
| total cholesterol | -1.302 | 16 | -1.110 | 16 | -2.930 | 16 | 1 | Clinical Chemistry | Physiology |
| total food intake | -1.938 | 6 | -1.755 | 6 | -3.002 | 6 | 1 | Indirect Calorimetry | Metabolism |
| total protein | -6.000 | 16 | -3.618 | 16 | -4.016 | 16 | 1 | Clinical Chemistry | Physiology |
| total water intake | -2.843 | 3 | -5.000 | 3 | -2.782 | 3 | 1 | Indirect Calorimetry | Metabolism |
| triglycerides | -2.116 | 16 | -1.701 | 16 | -1.947 | 16 | 1 | Clinical Chemistry | Physiology |
| urea (blood urea nitrogen - bun) | -1.658 | 16 | -1.442 | 16 | -2.669 | 16 | 1 | Clinical Chemistry | Physiology |
| uric acid | -6.000 | 3 | -1.838 | 3 | -1.085 | 3 | 1 | Clinical Chemistry | Physiology |
| white blood cell count | -2.181 | 16 | -1.698 | 16 | -2.411 | 16 | 1 | Hematology | Hematology |
| whole arena average speed | -2.253 | 13 | -2.232 | 13 | -2.518 | 13 | 1 | Open Field | Behaviour |
| whole arena permanence | -1.055 | 2 | -1.056 | 2 | -16.871 | 13 | 1 | Open Field | Behaviour |
| whole arena resting time | -2.730 | 13 | -2.866 | 13 | -2.432 | 13 | 1 | Open Field | Behaviour |

```
#### Meta-analysis of heterogeneity

## Perform meta-meta-analysis (3 for each of the 9 grouping terms: var.CVR, var.VR, var.RR)

metacombohetero_final <- metacombohetero %>%
  group_by(GroupingTerm) %>%
  nest()

# Final random effects meta-analyses within grouping terms, with SE of the estimate

heterog1 <- metacombohetero_final %>%

  mutate(
    model_heteroCVR = map(data, ~ metafor::rma.uni(
      yi = .x$var.CVR, sei = sqrt(1 / 2 * (.x$N.CVR - 1)),
      control = list(optimizer = "optim", 
                     optmethod = "Nelder-Mead",
                     maxit = 10000, 
                     stepadj = 0.5), verbose = F)),
    model_heteroVR = map(data, ~ metafor::rma.uni(
      yi = .x$var.VR, sei = sqrt(1 / 2 * (.x$N.VR - 1)),
      control = list(optimizer = "optim", 
                     optmethod = "Nelder-Mead", 
                     maxit = 10000, 
                     stepadj = 0.5), verbose = F)),
    model_heteroRR = map(data, ~ metafor::rma.uni(
      yi = .x$var.RR, sei = sqrt(1 / 2 * (.x$N.RR - 1)),
      control = list(optimizer = "optim", 
                     optmethod = "Nelder-Mead", 
                     maxit = 10000, 
                     stepadj = 0.5), verbose = F)) )

# Across all grouping terms   

metacombohetero_all_final <- metacombohetero %>%
  nest(data = everything()) 

# Final random effects meta-analyses ACROSS grouping terms, with SE of the estimate

heterog1_all <- metacombohetero_all_final %>%
  
  mutate( model_heteroCVR = map(data, ~ metafor::rma.uni(
      yi = .x$var.CVR, sei = sqrt(1 / 2 * (.x$N.CVR - 1)),
      control = list(optimizer = "optim", 
                     optmethod = "Nelder-Mead", 
                     maxit = 10000, 
                     stepadj = 0.5), verbose = F)),
    model_heteroVR = map(data, ~ metafor::rma.uni(yi = .x$var.VR, sei = sqrt(1 / 2 * (.x$N.VR - 1)),
      control = list(optimizer = "optim", 
                     optmethod = "Nelder-Mead", 
                     maxit = 10000, 
                     stepadj = 0.5), verbose = F)),
    model_heteroRR = map(data, ~ metafor::rma.uni(yi = .x$var.RR, sei = sqrt(1 / 2 * (.x$N.RR - 1)),
      control = list(optimizer = "optim", 
                     optmethod = "Nelder-Mead", 
                     maxit = 10000, 
                     stepadj = 0.5), verbose = F)))


# Re-structure data for each grouping term; extract heterogenenity/variance terms; delete un-used variables

extract_heterogeneity <- function(data, type){
  
  tmp <- data %>%
         filter(., GroupingTerm == type) %>%
         select(., -data) %>%
         mutate(heteroCVR       = .[[2]][[1]]$b, 
                heteroCVR_lower = .[[2]][[1]]$ci.lb, 
                heteroCVR_upper = .[[2]][[1]]$ci.ub, 
                heteroCVR_se    = .[[2]][[1]]$se,
                heteroVR        = .[[3]][[1]]$b, 
                heteroVR_lower  = .[[3]][[1]]$ci.lb, 
                heteroVR_upper  = .[[3]][[1]]$ci.ub, 
                heteroVR_se     = .[[3]][[1]]$se,
                heteroRR        = .[[4]][[1]]$b, 
                heteroRR_lower  = .[[4]][[1]]$ci.lb, 
                heteroRR_upper  = .[[4]][[1]]$ci.ub, 
                heteroRR_se     = .[[4]][[1]]$se) %>%
        select(., GroupingTerm, heteroCVR:heteroRR_se)
  
  return(tmp)
}

   Behaviour. <- extract_heterogeneity(heterog1, type = "Behaviour")
  Immunology. <- extract_heterogeneity(heterog1, type = "Immunology")
  Hematology. <- extract_heterogeneity(heterog1, type = "Hematology")
     Hearing. <- extract_heterogeneity(heterog1, type = "Hearing")
  Physiology. <- extract_heterogeneity(heterog1, type = "Physiology")
  Metabolism. <- extract_heterogeneity(heterog1, type = "Metabolism")
  Morphology. <- extract_heterogeneity(heterog1, type = "Morphology")
       Heart. <- extract_heterogeneity(heterog1, type = "Heart")
         Eye. <- extract_heterogeneity(heterog1, type = "Eye")

#Reorder to be able to keep cell referencing 
heterog1_all <- heterog1_all %>% 
                mutate(GroupingTerm = "All") %>% 
                select(GroupingTerm, everything())

All. <- heterog1_all %>% 
        select(., -data) %>%
        mutate(heteroCVR       = .[[2]][[1]]$b, 
               heteroCVR_lower = .[[2]][[1]]$ci.lb, 
               heteroCVR_upper = .[[2]][[1]]$ci.ub, 
               heteroCVR_se    = .[[2]][[1]]$se, 
               heteroVR        = .[[3]][[1]]$b, 
               heteroVR_lower  = .[[3]][[1]]$ci.lb, 
               heteroVR_upper  = .[[3]][[1]]$ci.ub, 
               heteroVR_se     = .[[3]][[1]]$se, 
               heteroRR        = .[[4]][[1]]$b, 
               heteroRR_lower  = .[[4]][[1]]$ci.lb, 
               heteroRR_upper  = .[[4]][[1]]$ci.ub, 
               heteroRR_se     = .[[4]][[1]]$se) %>%
        select(., GroupingTerm, heteroCVR:heteroRR_se)

heterog2 <- bind_rows(Behaviour., Morphology., Metabolism., Physiology., Immunology., Hematology., Heart., Hearing., Eye., All.)

#### Preparation of the Heterogeneity PLOT

#Restructure data for plotting

heterog3 <- gather(heterog2, parameter, value, c(heteroCVR, heteroVR, heteroRR), factor_key = TRUE)

heteroCVR.ci <- heterog3 %>%
                filter(parameter == "heteroCVR") %>%
                mutate(ci.low = heteroCVR_lower, 
                       ci.high = heteroCVR_upper)
heteroVR.ci <-  heterog3 %>%
                filter(parameter == "heteroVR") %>%
                mutate(ci.low = heteroVR_lower, 
                       ci.high = heteroVR_upper)
heteroRR.ci <-  heterog3 %>%
                filter(parameter == "heteroRR") %>%
                mutate(ci.low = heteroRR_lower, 
                       ci.high = heteroRR_upper)

heterog4 <- bind_rows(heteroCVR.ci, heteroVR.ci, heteroRR.ci) %>% select(GroupingTerm, parameter, value, ci.low, ci.high)

# **Re-order grouping terms

heterog4$GroupingTerm <- factor(heterog4$GroupingTerm, 
                                levels = c("Behaviour", "Morphology", "Metabolism", 
                                           "Physiology", "Immunology", "Hematology", "Heart", 
                                           "Hearing", "Eye", "All"))

heterog4$GroupingTerm <- factor(heterog4$GroupingTerm, 
                                rev(levels(heterog4$GroupingTerm)))
heterog4$label <- "All traits"
# write.csv(heterog4, "heterog4.csv")

#### Plot S1 C (Second-order meta analysis on heterogeneity)

heterog5 <- heterog4
heterog5$mean <- as.numeric(exp(heterog5$value))
heterog5$ci.l <- as.numeric(exp(heterog5$ci.low))
heterog5$ci.h <- as.numeric(exp(heterog5$ci.high))

heterog6 <- heterog5

HeteroS1 <-
  heterog6 %>%
  ggplot(aes(y = GroupingTerm, x = mean)) +
  geom_errorbarh(aes(
    xmin = ci.l,
    xmax = ci.h
  ),
  height = 0.1, show.legend = FALSE
  ) +
  geom_point(aes(shape = parameter),
    fill = "black",
    color = "black", size = 2.2,
    show.legend = FALSE
  ) +
  scale_x_continuous(
    limits = c(-0.1, 1.4),
    # breaks = c(0, 0.1, 0.2),
    name = "sigma^2"
  ) +
  # geom_vline(xintercept=0,
  # color='black',
  # linetype='dashed')+
  facet_grid(
    cols = vars(parameter), rows = vars(label),
    labeller = label_wrap_gen(width = 23),
    scales = "free",
    space = "free"
  ) +
  theme_bw() +
  theme(
    strip.text.y = element_text(angle = 270, size = 10, margin = margin(t = 15, r = 15, b = 15, l = 15)),
    strip.text.x = element_text(size = 12),
    strip.background = element_rect(colour = NULL, linetype = "blank", fill = "gray90"),
    text = element_text(size = 14),
    panel.spacing = unit(0.5, "lines"),
    panel.border = element_blank(),
    axis.line = element_line(),
    panel.grid.major.x = element_line(linetype = "solid", colour = "gray95"),
    panel.grid.major.y = element_line(linetype = "solid", color = "gray95"),
    panel.grid.minor.y = element_blank(),
    panel.grid.minor.x = element_blank(),
    legend.title = element_blank(),
    axis.title.x = element_text(hjust = 0.5, size = 14),
    axis.title.y = element_blank())
```

### 4 Meta-analysis results

#### 4.1 Preparing: Figure 3

##### 4.1.1 Count data, based on First-order meta-analysis results

This includes all separate eligible traits.
Re-ordering of grouping terms. Preparation of subplot 3A

```
meta_clean$GroupingTerm <- factor(meta_clean$GroupingTerm, 
                                  levels = c("Behaviour", "Morphology", "Metabolism", "Physiology", "Immunology", "Hematology", "Heart", "Hearing", "Eye"))
meta_clean$GroupingTerm <- factor(meta_clean$GroupingTerm, 
                                  rev(levels(meta_clean$GroupingTerm)))

# *Preparing data for all traits

meta.plot2.all <- meta_clean %>%
                  select(lnCVR, lnVR, lnRR, GroupingTerm) %>%
                  arrange(GroupingTerm)

# lnVR has been removed here and in the steps below, as this is only included in the supplemental figure
      meta.plot2.all.b <- gather(meta.plot2.all, trait, value, c(lnCVR, lnRR)) 

meta.plot2.all.b$trait <- factor(meta.plot2.all.b$trait, levels = c("lnCVR", "lnRR")) 
      meta.plot2.all.c <- meta.plot2.all.b %>%
                          group_by_at(vars(trait, GroupingTerm)) %>%
                          summarise(
                            malebias = sum(value > 0), femalebias = sum(value <= 0), total = malebias + femalebias,
                            malepercent = malebias * 100 / total, femalepercent = femalebias * 100 / total
                          )

meta.plot2.all.c$label <- "All traits"

# Re-structure to create stacked bar plots

meta.plot2.all.d <- as.data.frame(meta.plot2.all.c)
meta.plot2.all.e <- gather(meta.plot2.all.d, 
                           key = sex, 
                           value = percent, 
                           malepercent:femalepercent, 
                           factor_key = TRUE)

# Create new sample size variable

meta.plot2.all.e$samplesize <- with(meta.plot2.all.e, 
                                    ifelse(sex == "malepercent", 
                                           malebias, 
                                           femalebias))

# Add summary row ('All') and re-arrange rows into correct order for plotting (warnings about coercing 'id' into character vector are ok)

meta.plot2.all.f <- meta.plot2.all.e %>% 
                    group_by(trait, sex) %>% 
                      summarise(GroupingTerm = "All", 
                                malebias = sum(malebias),
                                femalebias = sum(femalebias),
                                total = malebias + femalebias, 
                                label = "All traits", 
                                samplesize = sum(samplesize)) %>%
                      mutate(percent = ifelse(sex == "femalepercent", femalebias*100/(malebias+femalebias), malebias*100/(malebias+femalebias))) %>%
                      bind_rows(meta.plot2.all.e, .) %>%
                      mutate(rownumber = row_number()) %>%
                      .[c(37, 1:9, 39, 10:18, 38, 19:27, 40, 28:36), ] 
  #line references in previous code line corresponding to: 
  #'lnCVR(male(All)), lnCVR(male('single grouping terms'), lnRR(male(All)), lnRR(male('single grouping terms')),
  #lnCVR(female(All)), lnCVR(female('single grouping terms'), lnRR(female(All)), lnRR(female('single grouping terms'))'

meta.plot2.all.f$GroupingTerm <- factor(meta.plot2.all.f$GroupingTerm,
                                        levels = c("Behaviour", "Morphology", "Metabolism", "Physiology", "Immunology", "Hematology", "Heart", "Hearing", "Eye", "All")) 
meta.plot2.all.f$GroupingTerm <- factor(meta.plot2.all.f$GroupingTerm, 
                                        rev(levels(meta.plot2.all.f$GroupingTerm)))

malebias_Fig2_alltraits <-
    ggplot(meta.plot2.all.f) +
    aes(x = GroupingTerm, y = percent, fill = sex) +
    ylab("Percentage") +
    geom_col() +
    geom_hline(yintercept = 50, linetype = "dashed", color = "gray40") +
    geom_text(
      data = subset(meta.plot2.all.f, samplesize != 0), aes(label = samplesize), position = position_stack(vjust = .5),
      color = "white", size = 3.5
    ) +
    facet_grid(
      cols = vars(trait), rows = vars(label), labeller = label_wrap_gen(width = 18),
      scales = "free", space = "free"
    ) +  
    scale_fill_brewer(palette = "Set2") +
    theme_bw(base_size = 18) +
    theme(
      strip.text.y = element_text(angle = 270, size = 10, margin = margin(t = 15, r = 15, b = 15, l = 15)),
      strip.text.x = element_text(size = 12),
      strip.background = element_rect(colour = NULL, linetype = "blank", fill = "gray90"),
      text = element_text(size = 14),
      panel.spacing = unit(0.5, "lines"),
      panel.border = element_blank(),
      axis.line = element_line(),
      panel.grid.major.x = element_line(linetype = "solid", colour = "gray95"),
      panel.grid.major.y = element_line(linetype = "solid", color = "gray95"),
      panel.grid.minor.y = element_blank(),
      panel.grid.minor.x = element_blank(),
      legend.position= "none",
      #axis.title.x = Percentage,
      axis.title.y = element_blank()
    ) +
    coord_flip()
```

##### 4.1.2 Re-structure data for plotting

Data are re-structured, and grouping terms are being re-ordered

```
overall3 <- gather(overall2, parameter, value, c(lnCVR, lnRR), factor_key = TRUE)

lnCVR.ci <- overall3 %>% filter(parameter == "lnCVR") %>% mutate(ci.low = lnCVR_lower, 
    ci.high = lnCVR_upper)
lnVR.ci <- overall3 %>% filter(parameter == "lnVR") %>% mutate(ci.low = lnVR_lower, 
    ci.high = lnVR_upper)
lnRR.ci <- overall3 %>% filter(parameter == "lnRR") %>% mutate(ci.low = lnRR_lower, 
    ci.high = lnRR_upper)

overall4 <- bind_rows(lnCVR.ci, lnRR.ci) %>% select(GroupingTerm, parameter, value, 
    ci.low, ci.high)

# Re-order grouping terms

overall4$GroupingTerm <- factor(overall4$GroupingTerm, levels = c("Behaviour", "Morphology", 
    "Metabolism", "Physiology", "Immunology", "Hematology", "Heart", "Hearing", "Eye", 
    "All"))
overall4$GroupingTerm <- factor(overall4$GroupingTerm, rev(levels(overall4$GroupingTerm)))
overall4$label <- "All traits"

#### PLOT 4B

# adding male / female symbols for Figure 4B additional packages for intergating
# male / female symbols in plot 4
library(png)
library(grid)

male <- readPNG(here("images", "MaleAqua.png"))
female <- readPNG(here("images", "FemaleSalmon.png"))


Metameta_Fig3_alltraits <- overall4 %>% 
ggplot(aes(y = GroupingTerm, x = value)) + geom_errorbarh(aes(xmin = ci.low, xmax = ci.high), 
    height = 0.1, show.legend = FALSE) + geom_point(aes(shape = parameter), fill = "black", 
    color = "black", size = 2.2, show.legend = FALSE) + scale_x_continuous(limits = c(-0.24, 
    0.25), breaks = c(-0.2, -0.1, 0, 0.1, 0.2), name = "Effect size") + geom_vline(xintercept = 0, 
    color = "black", linetype = "dashed") + facet_grid(cols = vars(parameter), rows = vars(label), 
    labeller = label_wrap_gen(width = 23), scales = "free", space = "free") + theme_bw() + 
    theme(strip.text.y = element_text(angle = 270, size = 10, margin = margin(t = 15, 
        r = 15, b = 15, l = 15)), strip.text.x = element_text(size = 12), strip.background = element_rect(colour = NULL, 
        linetype = "blank", fill = "gray90"), text = element_text(size = 14), panel.spacing = unit(0.5, 
        "lines"), panel.border = element_blank(), axis.line = element_line(), panel.grid.major.x = element_line(linetype = "solid", 
        colour = "gray95"), panel.grid.major.y = element_line(linetype = "solid", 
        color = "gray95"), panel.grid.minor.y = element_blank(), panel.grid.minor.x = element_blank(), 
        legend.title = element_blank(), axis.title.x = element_text(hjust = 0.5, 
            size = 14), axis.title.y = element_blank()) + # addition of male & female symols
annotation_custom(rasterGrob(female), xmin = -0.2, xmax = -0.1, ymin = 2.3, ymax = 4) + 
    annotation_custom(rasterGrob(male), xmin = 0.1, xmax = 0.2, ymin = 2.3, ymax = 4)

# Metameta_Fig3_alltraits
```

#### 4.2 Main Manuscript: Figure 3

```
Fig4 <- ggarrange(malebias_Fig2_alltraits, Metameta_Fig3_alltraits, nrow = 2, align = "v", 
    heights = c(10, 9), labels = c("A", "B"))
Fig4
```

Figure 4.1: Panel A shows the numbers of traits across functional groups that are either male-biased (blue-green) or female-biased (orange-red), as calculated in Figure 1.1D. Panel B shows effect sizes and 95% CI from separate meta-analysis for each functional group (step H in Figure1.1). Both panels represent results evaluated across all traits. Traits that are male biased are shown in blue, whereas female bias data is represented in orange.

### 5 Supplemental Figures

#### 5.1 Figure 5.1

Figure 5.1 is similar to Figure 3 in the main article except that lnVR has been added to the plot for comparison with lnCVR and lnRR.

```
#### Combined Figure S1: overall Count data, Meta anlysis results, Heterogeneity

FigS1 <- ggarrange(malebias_FigS1_alltraits, Metameta_FigS1_alltraits, HeteroS1, 
    nrow = 3, align = "v", heights = c(1.1, 1, 1), labels = c("A", "B", "C"))
FigS1
```

Figure 5.1: Numbers of either male (blue-green bars) or female (orange-red bars) biased traits (Panel A) across functional groups, this time for lnCVR (left hand side), lnVR (middle) and lnRR (right hand side). Panel B shows effect sizes from separate meta-analysis for each functional group, and Panel C contains results of heterogeneity analyses. All three panels represent results evaluated across all traits.

```
# ggsave('FigS1_OverallResults.pdf', plot = FigS1, width = 6, height = 7)
```

#### 5.2 Figure 5.2

##### 5.2.1 Sex-bias in traits sub-analysis

###### 5.2.1.1 Methods

Among the 147 traits (i.e. with ‘first-order’ meta-analytic results), we found 31 traits to be significantly male-biased and 30 female-biased for lnCVR (47 male-biased and 39 female- biased for lnRR; 74 male and 55 female-biased for lnVR). We used this statistical significance to categorize traits into two groups: male-based and female-biased traits. For each set of these sex-biased traits, we conducted second-order meta-analyses with trait as the unit of analysis in the same way as those using lnRR, lnVR and lnCVR (Figure 1.1J). As sensitivity analyses, we also employed another way of separating male- and female-biased traits. We quantified traits that were more than 10% biased towards either of the sexes, regardless of significance (N[male-biased] = 31, & N[female-biased] = 36 for lnCVR, N[male-biased] = 56, & N[female-biased] = 57 for lnVR , N[male-biased] = 42, & N[female-biased] = 33 for lnRR; see Supplemental Figure S5.2, Supplemental Table 7.3).

###### 5.2.1.2 Results

Having established that different sex-biases were present across trait functional groups, we explored specific patterns in male and female bias by categorizing traits into either male or female-biased, and then, quantifying the degree of sex bias. To do this, we focused on only those traits that were significantly biased towards one sex (as indicated by “CI not overlapping 0” in Figure S5.2. Selecting significantly sex-biased traits reduced the number of traits substantially (Figure S5.2A; compared to all traits, Figure 4.1 and Figure 5.1), and returned similar numbers of traits with either significant male- or female-bias for lnCVR (Figure S5.2A).

Looking only at traits with significant male-bias, trait variability tended to be higher in males than in females overall (11.77% male bias, lnCVR = 0.111, CI = [0.093 to 0.129]; compared to 8.3% female-bias, lnCVR = -0.087, CI = [-0.109 to (-0.064)]; Figure S5.2B). The degree of sex-bias varied across the functional trait groups, ranging from 4.6 % (Physiology, lnCVR = -0.047, CI = [-0.064 to (-0.030)] to 15.13 % (Eye morphology, lnCVR = -0.164, CI = [-0.249 to (-0.250)] in females, and from 8.6 % (Hematology, lnCVR = 0.082, CI = [0.048 to 0.117]) and 16.66 % (Heart, lnCVR = 0.154, CI = [0.085 to 0.223]) in males (Figure S5.2B).

To put sex biases (sexual dimorphisms) in variability into perspective, we also quantify the degree of sexual dimorphism in significantly sex-biased traits (i.e., the extent of sexual dimorphisms). There were more traits that were significantly biased in mean trait value in males compared to females (Figure S5.2A, lnRR). These traits were 11.73 % biased towards males (Figure S5.2B; lnRR = 0.111, CI = [0.083 – 0.138]) while 9.38 % towards females (lnRR = -0.094, CI = [-0.129 – (-0.068)]. The overall pattern for lnRR was more varied among the functional trait groups than for lnCVR, ranging from 1.2% (Hematology, lnRR = -0.013, CI = [-0.017 – (-0.007)] to 14.72 % (Immunology, lnRR = -0.159, CI = [-0.229 – (-0.089)] in female-based traits (Figure S5.2B) whereas in males-based traits from 1.8 % (Hearing, lnRR = 0.018, CI = [0.006 – 0.031]) to 25 % (Metabolism, lnRR = 0.223, CI = [0.110 – 0.335]) (Figure S5.2B).

```
library(ggpubr)
FigS2b <- ggarrange(Metameta_FigS2_female.sig, Metameta_FigS2_male.sig, ncol = 2, 
    nrow = 1, widths = c(1, 1.3), heights = c(1.2, 1.2))

FigS2d <- ggarrange(Metameta_Fig3_female.perc, Metameta_Fig3_male.perc, ncol = 2, 
    nrow = 1, widths = c(1, 1.2), heights = c(1.1, 1.1))

# end combination Figure 5
FigS2 <- ggarrange(malebias_FigS2_sigtraits, malebias_Fig2_over10, FigS2b, FigS2d, 
    ncol = 1, nrow = 4, heights = c(2.5, 2.2, 2.1, 2), labels = c("A", " ", "B", 
        " "))
FigS2
```

Figure 5.2: A) Differences in numbers of affected traits, in variance (lnCVR and lnVR) and means (lnRR), where there’s a significant difference between the sexes (i.e CI not overlapping zero), and where the sex bias is greater than 10% difference (regardless of significance). Panel B depicts results for the sex bias in those traits that differ between the sexes (second-order meta-analysis). Triangles represent sex bias in means (response ratio) and black circles differences in the coefficient of variation ratio (mean-adjusted variability). The orange-red bars represent trait groups with a female bias, blue-green bars male-biased traits.

### 6 Supplemental Tables

#### 6.1 Table 6.1

```
# PDF pander(ST1, caption = 'Table 6.1: Summary of the available numbers of male
# and female mice from each strain and originating institution.') HTML
kable(ST1, caption = "Summary of the available numbers of male and female mice from each strain and originating institution.") %>% 
    kable_styling() %>% scroll_box(width = "100%", height = "200px")
```

Table 6.1: Summary of the available numbers of male and female mice from each strain and originating institution.

| production\_center | strain\_name | sex | n |
| --- | --- | --- | --- |
| BCM | C57BL/6N | female | 653 |
| BCM | C57BL/6N | male | 639 |
| BCM | C57BL/6N;C57BL/6NTac | female | 47 |
| BCM | C57BL/6N;C57BL/6NTac | male | 52 |
| BCM | C57BL/6NCrl | female | 4 |
| BCM | C57BL/6NCrl | male | 2 |
| BCM | C57BL/6NJ | female | 6 |
| BCM | C57BL/6NJ | male | 6 |
| BCM | C57BL/6NTac | female | 1 |
| BCM | C57BL/6NTac | male | 5 |
| HMGU | C57BL/6NCrl | female | 313 |
| HMGU | C57BL/6NCrl | male | 311 |
| HMGU | C57BL/6NTac | female | 1045 |
| HMGU | C57BL/6NTac | male | 1062 |
| ICS | C57BL/6N | female | 1025 |
| ICS | C57BL/6N | male | 1050 |
| JAX | C57BL/6NJ | female | 2025 |
| JAX | C57BL/6NJ | male | 2022 |
| KMPC | C57BL/6N;C57BL/6NTac | female | 271 |
| KMPC | C57BL/6N;C57BL/6NTac | male | 266 |
| MARC | C57BL/6N | female | 936 |
| MARC | C57BL/6N | male | 926 |
| MRC Harwell | C57BL/6NTac | female | 2639 |
| MRC Harwell | C57BL/6NTac | male | 2661 |
| MRC Harwell | C57BL/6NTac | no\_data | 3 |
| RBRC | C57BL/6NJcl | female | 222 |
| RBRC | C57BL/6NJcl | male | 222 |
| RBRC | C57BL/6NTac | female | 526 |
| RBRC | C57BL/6NTac | male | 523 |
| TCP | C57BL/6NCrl | female | 552 |
| TCP | C57BL/6NCrl | male | 524 |
| TCP | C57BL6/NCrl | female | 2 |
| TCP | C57BL6/NCrl | male | 2 |
| UC Davis | C57BL/6N | male | 1 |
| UC Davis | C57BL/6NCrl | female | 1155 |
| UC Davis | C57BL/6NCrl | male | 1158 |
| WTSI | B6Brd;B6Dnk;B6N-Tyr<c-Brd> | female | 97 |
| WTSI | B6Brd;B6Dnk;B6N-Tyr<c-Brd> | male | 87 |
| WTSI | C57BL/6J-Tyr<c-Brd> or C57BL/6NTac/USA | male | 3 |
| WTSI | C57BL/6N | female | 1951 |
| WTSI | C57BL/6N | male | 2008 |
| WTSI | C57BL/6N;C57BL/6NTac | female | 41 |
| WTSI | C57BL/6N;C57BL/6NTac | male | 7 |
| WTSI | C57BL/6NCrl | male | 13 |
| WTSI | C57BL/6NTac | female | 49 |
| WTSI | C57BL/6NTac | male | 34 |

#### 6.2 Table 6.2

```
# Table detailing numbers of correlated and uncorrelated traits

# PDF pander(cbind(meta1 %>% count(parameter_group)), caption = 'Table 6.2: Trait
# categories (parameter_group) and the number of correlated traits within these
# categories. Traits were meta-analysed using robumeta')

kable(cbind(meta1 %>% count(parameter_group)), caption = "Trait categories (parameter_group) and the number of correlated traits within these categories. Traits were meta-analysed using robumeta") %>% 
    kable_styling() %>% scroll_box(width = "100%", height = "200px")
```

Table 6.2: Trait categories (parameter\_group) and the number of correlated traits within these categories. Traits were meta-analysed using robumeta

| parameter\_group | n |
| --- | --- |
| 12khz-evoked abr threshold | 1 |
| 18khz-evoked abr threshold | 1 |
| 24khz-evoked abr threshold | 1 |
| 30khz-evoked abr threshold | 1 |
| 6khz-evoked abr threshold | 1 |
| alanine aminotransferase | 1 |
| albumin | 1 |
| alkaline phosphatase | 1 |
| alpha-amylase | 1 |
| area under glucose response curve | 1 |
| aspartate aminotransferase | 1 |
| B cells | 4 |
| basophil cell count | 1 |
| basophil differential count | 1 |
| bmc/body weight | 1 |
| body length | 1 |
| body temp | 1 |
| body weight | 1 |
| body weight after experiment | 1 |
| body weight before experiment | 1 |
| bone area | 1 |
| bone mineral content (excluding skull) | 1 |
| bone mineral density (excluding skull) | 1 |
| calcium | 1 |
| cardiac output | 1 |
| cd4 nkt | 6 |
| cd4 t | 7 |
| cd8 nkt | 6 |
| cd8 t | 7 |
| cdcs | 2 |
| center average speed | 1 |
| center distance travelled | 1 |
| center permanence time | 1 |
| center resting time | 1 |
| chloride | 1 |
| click-evoked abr threshold | 1 |
| creatine kinase | 1 |
| creatinine | 1 |
| cv | 1 |
| distance travelled - total | 1 |
| dn nkt | 6 |
| dn t | 7 |
| ejection fraction | 1 |
| end-diastolic diameter | 1 |
| end-systolic diameter | 1 |
| eosinophils | 3 |
| fasted blood glucose concentration | 1 |
| fat mass | 1 |
| fat/body weight | 1 |
| follicular b cells | 2 |
| forelimb and hindlimb grip strength measurement mean | 1 |
| forelimb grip strength measurement mean | 1 |
| fractional shortening | 1 |
| free fatty acids | 1 |
| fructosamine | 1 |
| glucose | 1 |
| hdl-cholesterol | 1 |
| heart weight | 1 |
| heart weight normalised against body weight | 1 |
| hematocrit | 1 |
| hemoglobin | 1 |
| hr | 1 |
| hrv | 1 |
| initial response to glucose challenge | 1 |
| insulin | 1 |
| iron | 1 |
| lactate dehydrogenase | 1 |
| latency to center entry | 1 |
| ldl-cholesterol | 1 |
| lean mass | 1 |
| lean/body weight | 1 |
| left anterior chamber depth | 1 |
| left corneal thickness | 1 |
| left inner nuclear layer | 1 |
| left outer nuclear layer | 1 |
| left posterior chamber depth | 1 |
| left total retinal thickness | 1 |
| locomotor activity | 1 |
| luc | 2 |
| lvawd | 1 |
| lvaws | 1 |
| lvidd | 1 |
| lvids | 1 |
| lvpwd | 1 |
| lvpws | 1 |
| lymphocytes | 2 |
| magnesium | 1 |
| mean cell hemoglobin concentration | 1 |
| mean cell volume | 1 |
| mean corpuscular hemoglobin | 1 |
| mean platelet volume | 1 |
| mean r amplitude | 1 |
| mean sr amplitude | 1 |
| monocytes | 3 |
| neutrophils | 3 |
| nk cells | 6 |
| nkt cells | 4 |
| number of center entries | 1 |
| number of rears - total | 1 |
| others | 1 |
| pdcs | 1 |
| percentage center time | 1 |
| percentage of live gated events | 2 |
| periphery average speed | 1 |
| periphery distance travelled | 1 |
| periphery permanence time | 1 |
| periphery resting time | 1 |
| phosphorus | 1 |
| platelet count | 1 |
| pnn5(6>ms) | 1 |
| potassium | 1 |
| pq | 1 |
| pr | 1 |
| pre-pulse inhibition | 5 |
| qrs | 1 |
| qtc | 1 |
| qtc dispersion | 1 |
| red blood cell count | 1 |
| red blood cell distribution width | 1 |
| respiration rate | 1 |
| respiratory exchange ratio | 1 |
| response amplitude | 10 |
| right anterior chamber depth | 1 |
| right corneal thickness | 1 |
| right inner nuclear layer | 1 |
| right outer nuclear layer | 1 |
| right posterior chamber depth | 1 |
| right total retinal thickness | 1 |
| rmssd | 1 |
| rp macrophage (cd19- cd11c-) | 1 |
| rr | 1 |
| sodium | 1 |
| spleen weight | 1 |
| st | 1 |
| stroke volume | 1 |
| t cells | 3 |
| tibia length | 1 |
| total bilirubin | 1 |
| total cholesterol | 1 |
| total food intake | 1 |
| total protein | 1 |
| total water intake | 1 |
| triglycerides | 1 |
| urea (blood urea nitrogen - bun) | 1 |
| uric acid | 1 |
| white blood cell count | 1 |
| whole arena average speed | 1 |
| whole arena resting time | 1 |

#### 6.3 Table 6.3

```
# PDF pander(metacombo, caption = 'Table 6.3: We use this corrected (for
# correlated traits) results table, which contains each of the meta-analytic
# means for all effect sizes of interest, for further analyses.  We further use
# this table as part of the Shiny App, which is able to provide the percentage
# differences between males and females for mean, variance and coefficient of
# variance.')

# HTML
kable(metacombo, digits = 3, caption = "We use this corrected (for correlated traits) results table, which contains each of the meta-analytic means for all effect sizes of interest, for further analyses.  We further use this table as part of the Shiny App, which is able to provide the percentage differences between males and females for mean, variance and coefficient of variance.") %>% 
    kable_styling() %>% scroll_box(width = "100%", height = "200px")
```

Table 6.3: We use this corrected (for correlated traits) results table, which contains each of the meta-analytic means for all effect sizes of interest, for further analyses. We further use this table as part of the Shiny App, which is able to provide the percentage differences between males and females for mean, variance and coefficient of variance.

| parameter\_group | counts | procedure | GroupingTerm | lnCVR | lnCVR\_lower | lnCVR\_upper | lnCVR\_se | lnVR | lnVR\_lower | lnVR\_upper | lnVR\_se | lnRR | lnRR\_lower | lnRR\_upper | lnRR\_se |
| --- | --- | --- | --- | --- | --- | --- | --- | --- | --- | --- | --- | --- | --- | --- | --- |
| pre-pulse inhibition | 5 | Acoustic Startle and Pre-pulse Inhibition (PPI) | Behaviour | 0.023 | -0.080 | 0.127 | 0.037 | 0.009 | -0.036 | 0.055 | 0.014 | -0.005 | -0.043 | 0.032 | 0.013 |
| B cells | 4 | Immunophenotyping | Immunology | -0.094 | -0.250 | 0.062 | 0.043 | -0.100 | -0.207 | 0.008 | 0.025 | -0.003 | -0.130 | 0.125 | 0.039 |
| cd4 nkt | 6 | Immunophenotyping | Immunology | -0.029 | -0.057 | -0.001 | 0.010 | -0.202 | -0.310 | -0.094 | 0.033 | -0.234 | -0.401 | -0.068 | 0.063 |
| cd4 t | 7 | Immunophenotyping | Immunology | -0.151 | -0.243 | -0.059 | 0.036 | -0.170 | -0.263 | -0.077 | 0.035 | -0.003 | -0.041 | 0.035 | 0.015 |
| cd8 nkt | 6 | Immunophenotyping | Immunology | -0.042 | -0.078 | -0.007 | 0.012 | -0.030 | -0.182 | 0.122 | 0.053 | 0.004 | -0.057 | 0.064 | 0.021 |
| cd8 t | 7 | Immunophenotyping | Immunology | -0.122 | -0.218 | -0.027 | 0.036 | -0.158 | -0.234 | -0.082 | 0.027 | -0.042 | -0.051 | -0.032 | 0.002 |
| cdcs | 2 | Immunophenotyping | Immunology | -0.036 | -0.359 | 0.286 | 0.025 | 0.108 | -0.057 | 0.273 | 0.013 | 0.164 | -0.170 | 0.499 | 0.026 |
| dn nkt | 6 | Immunophenotyping | Immunology | -0.062 | -0.136 | 0.012 | 0.026 | -0.157 | -0.281 | -0.033 | 0.045 | -0.173 | -0.291 | -0.055 | 0.044 |
| dn t | 7 | Immunophenotyping | Immunology | -0.080 | -0.184 | 0.025 | 0.042 | -0.242 | -0.343 | -0.141 | 0.041 | -0.230 | -0.252 | -0.208 | 0.007 |
| eosinophils | 3 | Hematology | Hematology | -0.066 | -0.281 | 0.148 | 0.033 | -0.015 | -0.405 | 0.374 | 0.087 | -0.004 | -0.241 | 0.232 | 0.051 |
| follicular b cells | 2 | Immunophenotyping | Immunology | -0.116 | -0.726 | 0.494 | 0.048 | -0.105 | -0.695 | 0.485 | 0.046 | 0.005 | -0.187 | 0.198 | 0.015 |
| luc | 2 | Hematology | Hematology | 0.018 | -0.204 | 0.240 | 0.017 | 0.266 | -1.225 | 1.757 | 0.117 | 0.222 | -1.414 | 1.857 | 0.129 |
| lymphocytes | 2 | Hematology | Hematology | 0.081 | -2.262 | 2.423 | 0.184 | 0.155 | -1.089 | 1.399 | 0.098 | 0.060 | -1.013 | 1.134 | 0.084 |
| monocytes | 3 | Hematology | Hematology | -0.021 | -0.203 | 0.160 | 0.042 | 0.078 | -0.181 | 0.338 | 0.059 | 0.103 | -0.148 | 0.353 | 0.057 |
| neutrophils | 3 | Hematology | Hematology | 0.259 | 0.013 | 0.504 | 0.056 | 0.380 | -0.206 | 0.966 | 0.132 | 0.132 | -0.267 | 0.531 | 0.092 |
| nk cells | 6 | Immunophenotyping | Immunology | -0.041 | -0.096 | 0.013 | 0.020 | 0.016 | -0.070 | 0.102 | 0.032 | 0.047 | -0.016 | 0.111 | 0.023 |
| nkt cells | 4 | Immunophenotyping | Immunology | 0.003 | -0.107 | 0.114 | 0.029 | -0.246 | -0.445 | -0.047 | 0.043 | -0.182 | -0.323 | -0.041 | 0.031 |
| percentage of live gated events | 2 | Immunophenotyping | Immunology | -0.093 | -0.304 | 0.117 | 0.017 | -0.041 | -0.141 | 0.059 | 0.008 | 0.050 | 0.008 | 0.092 | 0.003 |
| response amplitude | 10 | Acoustic Startle and Pre-pulse Inhibition (PPI) | Behaviour | 0.033 | -0.013 | 0.079 | 0.020 | 0.255 | 0.197 | 0.313 | 0.026 | 0.202 | 0.111 | 0.292 | 0.040 |
| t cells | 3 | Immunophenotyping | Immunology | -0.134 | -0.275 | 0.007 | 0.033 | -0.124 | -0.412 | 0.164 | 0.067 | -0.001 | -0.166 | 0.165 | 0.037 |
| 12khz-evoked abr threshold | 1 | Auditory Brain Stem Response | Hearing | 0.054 | -0.006 | 0.113 | 0.030 | 0.087 | 0.007 | 0.167 | 0.041 | 0.002 | -0.021 | 0.026 | 0.012 |
| 18khz-evoked abr threshold | 1 | Auditory Brain Stem Response | Hearing | 0.024 | -0.033 | 0.081 | 0.029 | 0.025 | -0.049 | 0.099 | 0.038 | -0.020 | -0.043 | 0.003 | 0.012 |
| 24khz-evoked abr threshold | 1 | Auditory Brain Stem Response | Hearing | 0.052 | -0.015 | 0.118 | 0.034 | -0.089 | -0.332 | 0.154 | 0.124 | -0.022 | -0.044 | 0.000 | 0.011 |
| 30khz-evoked abr threshold | 1 | Auditory Brain Stem Response | Hearing | 0.017 | -0.053 | 0.088 | 0.036 | -0.034 | -0.102 | 0.033 | 0.034 | -0.050 | -0.075 | -0.025 | 0.013 |
| 6khz-evoked abr threshold | 1 | Auditory Brain Stem Response | Hearing | -0.008 | -0.042 | 0.026 | 0.017 | 0.014 | -0.019 | 0.047 | 0.017 | 0.018 | 0.006 | 0.031 | 0.006 |
| alanine aminotransferase | 1 | Clinical Chemistry | Physiology | -0.068 | -0.190 | 0.053 | 0.062 | 0.059 | -0.132 | 0.249 | 0.097 | 0.107 | 0.032 | 0.182 | 0.038 |
| albumin | 1 | Clinical Chemistry | Physiology | 0.113 | 0.045 | 0.181 | 0.035 | 0.056 | -0.008 | 0.120 | 0.033 | -0.057 | -0.073 | -0.040 | 0.008 |
| alkaline phosphatase | 1 | Clinical Chemistry | Physiology | 0.104 | 0.045 | 0.164 | 0.030 | -0.311 | -0.398 | -0.224 | 0.044 | -0.422 | -0.469 | -0.374 | 0.024 |
| alpha-amylase | 1 | Clinical Chemistry | Physiology | 0.038 | -0.042 | 0.119 | 0.041 | 0.280 | 0.162 | 0.398 | 0.060 | 0.225 | 0.179 | 0.270 | 0.023 |
| area under glucose response curve | 1 | Intraperitoneal glucose tolerance test (IPGTT) | Metabolism | -0.153 | -0.221 | -0.085 | 0.035 | 0.275 | 0.195 | 0.355 | 0.041 | 0.436 | 0.366 | 0.506 | 0.036 |
| aspartate aminotransferase | 1 | Clinical Chemistry | Physiology | 0.012 | -0.123 | 0.147 | 0.069 | -0.057 | -0.246 | 0.132 | 0.096 | -0.059 | -0.133 | 0.016 | 0.038 |
| basophil cell count | 1 | Hematology | Hematology | -0.092 | -0.202 | 0.019 | 0.056 | 0.203 | -0.013 | 0.419 | 0.110 | 0.268 | 0.064 | 0.471 | 0.104 |
| basophil differential count | 1 | Hematology | Hematology | -0.093 | -0.179 | -0.008 | 0.044 | -0.064 | -0.283 | 0.155 | 0.112 | -0.016 | -0.110 | 0.079 | 0.048 |
| bmc/body weight | 1 | Body Composition (DEXA lean/fat) | Morphology | 0.131 | 0.033 | 0.230 | 0.050 | -0.045 | -0.134 | 0.044 | 0.045 | -0.172 | -0.221 | -0.124 | 0.025 |
| body length | 1 | Body Composition (DEXA lean/fat) | Morphology | -0.035 | -0.082 | 0.013 | 0.024 | -0.006 | -0.053 | 0.041 | 0.024 | 0.028 | 0.023 | 0.033 | 0.003 |
| body temp | 1 | Echo | Heart | -0.033 | -0.107 | 0.042 | 0.038 | -0.030 | -0.104 | 0.044 | 0.038 | 0.002 | -0.001 | 0.004 | 0.001 |
| body weight | 1 | Body Weight | Morphology | 0.025 | -0.042 | 0.091 | 0.034 | 0.234 | 0.169 | 0.298 | 0.033 | 0.210 | 0.194 | 0.225 | 0.008 |
| body weight after experiment | 1 | Indirect Calorimetry | Metabolism | 0.085 | 0.030 | 0.141 | 0.028 | 0.285 | 0.233 | 0.337 | 0.027 | 0.203 | 0.186 | 0.220 | 0.009 |
| body weight before experiment | 1 | Indirect Calorimetry | Metabolism | 0.105 | 0.041 | 0.169 | 0.033 | 0.304 | 0.244 | 0.364 | 0.031 | 0.201 | 0.182 | 0.220 | 0.010 |
| bone area | 1 | Body Composition (DEXA lean/fat) | Morphology | 0.098 | 0.027 | 0.169 | 0.036 | 0.129 | 0.053 | 0.204 | 0.038 | 0.032 | 0.000 | 0.063 | 0.016 |
| bone mineral content (excluding skull) | 1 | Body Composition (DEXA lean/fat) | Morphology | 0.171 | 0.063 | 0.279 | 0.055 | 0.209 | 0.102 | 0.317 | 0.055 | 0.037 | -0.013 | 0.088 | 0.026 |
| bone mineral density (excluding skull) | 1 | Body Composition (DEXA lean/fat) | Morphology | 0.054 | -0.088 | 0.197 | 0.073 | 0.049 | -0.109 | 0.207 | 0.081 | 0.001 | -0.019 | 0.021 | 0.010 |
| calcium | 1 | Clinical Chemistry | Physiology | 0.010 | -0.046 | 0.066 | 0.029 | 0.014 | -0.042 | 0.070 | 0.029 | 0.004 | 0.000 | 0.007 | 0.002 |
| cardiac output | 1 | Echo | Heart | 0.013 | -0.080 | 0.107 | 0.048 | 0.102 | 0.021 | 0.183 | 0.041 | 0.093 | 0.058 | 0.129 | 0.018 |
| center average speed | 1 | Open Field | Behaviour | 0.017 | -0.040 | 0.074 | 0.029 | -0.059 | -0.100 | -0.017 | 0.021 | -0.072 | -0.115 | -0.030 | 0.022 |
| center distance travelled | 1 | Open Field | Behaviour | -0.016 | -0.073 | 0.041 | 0.029 | -0.106 | -0.202 | -0.010 | 0.049 | -0.094 | -0.195 | 0.007 | 0.051 |
| center permanence time | 1 | Open Field | Behaviour | -0.025 | -0.083 | 0.032 | 0.029 | -0.026 | -0.101 | 0.050 | 0.039 | -0.004 | -0.090 | 0.083 | 0.044 |
| center resting time | 1 | Open Field | Behaviour | 0.024 | -0.074 | 0.123 | 0.050 | -0.023 | -0.155 | 0.109 | 0.067 | -0.063 | -0.222 | 0.095 | 0.081 |
| chloride | 1 | Clinical Chemistry | Physiology | 0.032 | -0.127 | 0.191 | 0.081 | 0.024 | -0.144 | 0.192 | 0.086 | -0.013 | -0.018 | -0.008 | 0.003 |
| click-evoked abr threshold | 1 | Auditory Brain Stem Response | Hearing | -0.053 | -0.153 | 0.048 | 0.051 | -0.056 | -0.183 | 0.071 | 0.065 | -0.015 | -0.058 | 0.027 | 0.022 |
| creatine kinase | 1 | Clinical Chemistry | Physiology | 0.024 | -0.107 | 0.155 | 0.067 | -0.132 | -0.397 | 0.133 | 0.135 | -0.134 | -0.384 | 0.115 | 0.127 |
| creatinine | 1 | Clinical Chemistry | Physiology | 0.035 | -0.023 | 0.093 | 0.030 | 0.107 | -0.220 | 0.433 | 0.167 | -0.084 | -0.132 | -0.037 | 0.024 |
| cv | 1 | Electrocardiogram (ECG) | Heart | 0.187 | 0.072 | 0.303 | 0.059 | -0.090 | -0.248 | 0.069 | 0.081 | -0.240 | -0.341 | -0.139 | 0.051 |
| distance travelled - total | 1 | Open Field | Behaviour | -0.019 | -0.086 | 0.048 | 0.034 | -0.127 | -0.200 | -0.055 | 0.037 | -0.112 | -0.182 | -0.043 | 0.035 |
| ejection fraction | 1 | Echo | Heart | -0.030 | -0.135 | 0.074 | 0.053 | -0.053 | -0.148 | 0.043 | 0.049 | -0.028 | -0.049 | -0.008 | 0.011 |
| end-diastolic diameter | 1 | Echo | Heart | 0.112 | 0.043 | 0.181 | 0.035 | 0.174 | 0.088 | 0.261 | 0.044 | 0.060 | 0.035 | 0.085 | 0.013 |
| end-systolic diameter | 1 | Echo | Heart | -0.008 | -0.078 | 0.061 | 0.036 | 0.067 | -0.002 | 0.135 | 0.035 | 0.076 | 0.045 | 0.108 | 0.016 |
| fasted blood glucose concentration | 1 | Intraperitoneal glucose tolerance test (IPGTT) | Metabolism | -0.018 | -0.126 | 0.090 | 0.055 | 0.070 | -0.030 | 0.171 | 0.051 | 0.087 | 0.049 | 0.124 | 0.019 |
| fat mass | 1 | Body Composition (DEXA lean/fat) | Morphology | 0.041 | -0.043 | 0.125 | 0.043 | 0.371 | 0.270 | 0.473 | 0.052 | 0.328 | 0.267 | 0.390 | 0.031 |
| fat/body weight | 1 | Body Composition (DEXA lean/fat) | Morphology | 0.078 | -0.012 | 0.167 | 0.046 | 0.202 | 0.108 | 0.296 | 0.048 | 0.124 | 0.064 | 0.183 | 0.030 |
| forelimb and hindlimb grip strength measurement mean | 1 | Grip Strength | Morphology | 0.058 | 0.004 | 0.112 | 0.027 | 0.115 | 0.053 | 0.176 | 0.031 | 0.054 | 0.029 | 0.079 | 0.013 |
| forelimb grip strength measurement mean | 1 | Grip Strength | Morphology | 0.027 | -0.019 | 0.072 | 0.023 | 0.100 | 0.054 | 0.145 | 0.023 | 0.070 | 0.044 | 0.096 | 0.013 |
| fractional shortening | 1 | Echo | Heart | -0.015 | -0.116 | 0.086 | 0.052 | -0.058 | -0.156 | 0.041 | 0.050 | -0.041 | -0.057 | -0.026 | 0.008 |
| free fatty acids | 1 | Clinical Chemistry | Physiology | 0.028 | -0.100 | 0.157 | 0.066 | 0.055 | -0.074 | 0.185 | 0.066 | 0.019 | -0.009 | 0.048 | 0.015 |
| fructosamine | 1 | Clinical Chemistry | Physiology | -0.040 | -0.120 | 0.040 | 0.041 | -0.068 | -0.151 | 0.016 | 0.043 | -0.028 | -0.069 | 0.013 | 0.021 |
| glucose | 1 | Clinical Chemistry | Physiology | 0.069 | 0.018 | 0.120 | 0.026 | 0.128 | 0.042 | 0.214 | 0.044 | 0.065 | 0.022 | 0.108 | 0.022 |
| hdl-cholesterol | 1 | Clinical Chemistry | Physiology | -0.065 | -0.126 | -0.004 | 0.031 | 0.172 | 0.070 | 0.275 | 0.052 | 0.261 | 0.218 | 0.303 | 0.022 |
| heart weight | 1 | Heart Weight | Morphology | 0.177 | 0.067 | 0.286 | 0.056 | 0.365 | 0.217 | 0.513 | 0.076 | 0.174 | 0.141 | 0.207 | 0.017 |
| heart weight normalised against body weight | 1 | Heart Weight | Morphology | 0.079 | -0.006 | 0.165 | 0.044 | 0.036 | -0.097 | 0.168 | 0.068 | -0.050 | -0.084 | -0.016 | 0.017 |
| hematocrit | 1 | Hematology | Hematology | 0.057 | -0.052 | 0.165 | 0.055 | 0.074 | -0.033 | 0.180 | 0.054 | 0.017 | 0.004 | 0.031 | 0.007 |
| hemoglobin | 1 | Hematology | Hematology | 0.087 | 0.027 | 0.146 | 0.030 | 0.087 | 0.019 | 0.154 | 0.034 | 0.005 | -0.008 | 0.018 | 0.007 |
| hr | 1 | Electrocardiogram (ECG) | Heart | -0.063 | -0.173 | 0.047 | 0.056 | -0.014 | -0.149 | 0.121 | 0.069 | 0.041 | -0.014 | 0.095 | 0.028 |
| hrv | 1 | Electrocardiogram (ECG) | Heart | 0.172 | 0.109 | 0.235 | 0.032 | -0.081 | -0.213 | 0.050 | 0.067 | -0.250 | -0.366 | -0.135 | 0.059 |
| initial response to glucose challenge | 1 | Intraperitoneal glucose tolerance test (IPGTT) | Metabolism | -0.097 | -0.150 | -0.043 | 0.027 | 0.043 | 0.014 | 0.072 | 0.015 | 0.118 | 0.085 | 0.151 | 0.017 |
| insulin | 1 | Insulin Blood Level | Metabolism | -0.099 | -0.372 | 0.174 | 0.139 | 0.177 | -0.194 | 0.549 | 0.189 | 0.445 | 0.094 | 0.795 | 0.179 |
| iron | 1 | Clinical Chemistry | Physiology | -0.097 | -0.214 | 0.019 | 0.060 | -0.253 | -0.396 | -0.111 | 0.073 | -0.153 | -0.193 | -0.113 | 0.021 |
| lactate dehydrogenase | 1 | Clinical Chemistry | Physiology | 0.094 | -0.021 | 0.210 | 0.059 | 0.141 | -0.062 | 0.344 | 0.104 | 0.032 | -0.141 | 0.205 | 0.088 |
| latency to center entry | 1 | Open Field | Behaviour | 0.125 | 0.033 | 0.218 | 0.047 | 0.364 | 0.206 | 0.523 | 0.081 | 0.273 | 0.074 | 0.473 | 0.102 |
| ldl-cholesterol | 1 | Clinical Chemistry | Physiology | 0.423 | 0.155 | 0.691 | 0.137 | 0.267 | -0.096 | 0.630 | 0.185 | -0.162 | -0.601 | 0.278 | 0.224 |
| lean mass | 1 | Body Composition (DEXA lean/fat) | Morphology | 0.144 | 0.076 | 0.211 | 0.035 | 0.338 | 0.266 | 0.410 | 0.037 | 0.193 | 0.175 | 0.211 | 0.009 |
| lean/body weight | 1 | Body Composition (DEXA lean/fat) | Morphology | 0.195 | 0.091 | 0.300 | 0.053 | 0.184 | 0.086 | 0.282 | 0.050 | -0.012 | -0.026 | 0.001 | 0.007 |
| left anterior chamber depth | 1 | Eye Morphology | Eye | -0.185 | -0.431 | 0.060 | 0.125 | -0.153 | -0.401 | 0.094 | 0.126 | 0.033 | 0.028 | 0.038 | 0.002 |
| left corneal thickness | 1 | Eye Morphology | Eye | -0.145 | -0.234 | -0.055 | 0.046 | -0.135 | -0.223 | -0.047 | 0.045 | 0.008 | -0.006 | 0.021 | 0.007 |
| left inner nuclear layer | 1 | Eye Morphology | Eye | 0.048 | -0.036 | 0.132 | 0.043 | 0.049 | -0.035 | 0.132 | 0.043 | 0.001 | -0.010 | 0.011 | 0.005 |
| left outer nuclear layer | 1 | Eye Morphology | Eye | -0.068 | -0.151 | 0.016 | 0.043 | -0.062 | -0.145 | 0.022 | 0.043 | 0.006 | 0.001 | 0.012 | 0.003 |
| left posterior chamber depth | 1 | Eye Morphology | Eye | -0.263 | -0.473 | -0.053 | 0.107 | -0.269 | -0.479 | -0.058 | 0.107 | -0.003 | -0.015 | 0.009 | 0.006 |
| left total retinal thickness | 1 | Eye Morphology | Eye | -0.198 | -0.439 | 0.044 | 0.123 | -0.193 | -0.427 | 0.040 | 0.119 | 0.003 | -0.003 | 0.009 | 0.003 |
| locomotor activity | 1 | Combined SHIRPA and Dysmorphology | Behaviour | 0.096 | 0.022 | 0.170 | 0.038 | -0.016 | -0.058 | 0.026 | 0.021 | -0.111 | -0.176 | -0.045 | 0.033 |
| lvawd | 1 | Echo | Heart | 0.023 | -0.025 | 0.070 | 0.024 | 0.045 | -0.001 | 0.092 | 0.024 | 0.025 | 0.011 | 0.038 | 0.007 |
| lvaws | 1 | Echo | Heart | -0.002 | -0.252 | 0.248 | 0.128 | 0.023 | -0.178 | 0.224 | 0.103 | 0.011 | -0.031 | 0.053 | 0.021 |
| lvidd | 1 | Echo | Heart | 0.045 | -0.024 | 0.115 | 0.035 | 0.098 | 0.021 | 0.175 | 0.039 | 0.053 | 0.038 | 0.068 | 0.008 |
| lvids | 1 | Echo | Heart | -0.064 | -0.199 | 0.072 | 0.069 | 0.008 | -0.134 | 0.150 | 0.072 | 0.076 | 0.053 | 0.099 | 0.012 |
| lvpwd | 1 | Echo | Heart | -0.032 | -0.126 | 0.062 | 0.048 | -0.010 | -0.127 | 0.106 | 0.060 | 0.030 | 0.013 | 0.047 | 0.009 |
| lvpws | 1 | Echo | Heart | -0.019 | -0.101 | 0.063 | 0.042 | 0.009 | -0.082 | 0.100 | 0.047 | 0.027 | 0.006 | 0.047 | 0.010 |
| magnesium | 1 | Urinalysis | Physiology | 0.016 | -0.023 | 0.055 | 0.020 | -0.051 | -0.117 | 0.014 | 0.033 | -0.041 | -0.114 | 0.031 | 0.037 |
| mean cell hemoglobin concentration | 1 | Hematology | Hematology | 0.038 | -0.088 | 0.164 | 0.064 | 0.025 | -0.109 | 0.159 | 0.068 | -0.011 | -0.015 | -0.008 | 0.002 |
| mean cell volume | 1 | Hematology | Hematology | 0.004 | -0.096 | 0.104 | 0.051 | -0.003 | -0.096 | 0.090 | 0.048 | -0.006 | -0.010 | -0.003 | 0.002 |
| mean corpuscular hemoglobin | 1 | Hematology | Hematology | -0.003 | -0.065 | 0.060 | 0.032 | -0.019 | -0.082 | 0.044 | 0.032 | -0.017 | -0.020 | -0.014 | 0.001 |
| mean platelet volume | 1 | Hematology | Hematology | 0.049 | -0.004 | 0.102 | 0.027 | 0.035 | -0.021 | 0.092 | 0.029 | -0.017 | -0.028 | -0.007 | 0.005 |
| mean r amplitude | 1 | Electrocardiogram (ECG) | Heart | 0.008 | -0.028 | 0.045 | 0.019 | -0.095 | -0.163 | -0.027 | 0.035 | -0.084 | -0.150 | -0.017 | 0.034 |
| mean sr amplitude | 1 | Electrocardiogram (ECG) | Heart | 0.028 | -0.013 | 0.070 | 0.021 | -0.088 | -0.127 | -0.048 | 0.020 | -0.113 | -0.156 | -0.070 | 0.022 |
| number of center entries | 1 | Open Field | Behaviour | 0.015 | -0.053 | 0.084 | 0.035 | -0.036 | -0.095 | 0.023 | 0.030 | -0.059 | -0.168 | 0.050 | 0.056 |
| number of rears - total | 1 | Open Field | Behaviour | -0.001 | -0.114 | 0.112 | 0.058 | 0.187 | -0.039 | 0.413 | 0.115 | 0.179 | 0.057 | 0.302 | 0.063 |
| others | 1 | Immunophenotyping | Immunology | -0.168 | -0.260 | -0.077 | 0.047 | -0.152 | -0.244 | -0.059 | 0.047 | 0.020 | 0.005 | 0.034 | 0.007 |
| pdcs | 1 | Immunophenotyping | Immunology | -0.173 | -0.400 | 0.054 | 0.116 | -0.257 | -0.719 | 0.204 | 0.235 | -0.092 | -0.252 | 0.069 | 0.082 |
| percentage center time | 1 | Open Field | Behaviour | -0.022 | -0.086 | 0.042 | 0.033 | -0.019 | -0.091 | 0.053 | 0.037 | -0.006 | -0.097 | 0.085 | 0.046 |
| periphery average speed | 1 | Open Field | Behaviour | -0.044 | -0.108 | 0.019 | 0.033 | -0.140 | -0.212 | -0.068 | 0.037 | -0.096 | -0.145 | -0.048 | 0.025 |
| periphery distance travelled | 1 | Open Field | Behaviour | -0.031 | -0.092 | 0.029 | 0.031 | -0.134 | -0.187 | -0.081 | 0.027 | -0.104 | -0.171 | -0.036 | 0.035 |
| periphery permanence time | 1 | Open Field | Behaviour | -0.037 | -0.128 | 0.054 | 0.046 | -0.029 | -0.101 | 0.042 | 0.036 | 0.008 | -0.014 | 0.029 | 0.011 |
| periphery resting time | 1 | Open Field | Behaviour | -0.054 | -0.127 | 0.019 | 0.037 | -0.057 | -0.107 | -0.007 | 0.025 | 0.003 | -0.056 | 0.061 | 0.030 |
| phosphorus | 1 | Clinical Chemistry | Physiology | -0.049 | -0.084 | -0.013 | 0.018 | -0.083 | -0.158 | -0.008 | 0.038 | -0.042 | -0.081 | -0.003 | 0.020 |
| platelet count | 1 | Hematology | Hematology | 0.074 | 0.021 | 0.127 | 0.027 | 0.242 | 0.187 | 0.296 | 0.028 | 0.164 | 0.137 | 0.191 | 0.014 |
| pnn5(6>ms) | 1 | Electrocardiogram (ECG) | Heart | 0.291 | 0.172 | 0.410 | 0.061 | -0.293 | -0.527 | -0.058 | 0.120 | -0.600 | -0.924 | -0.277 | 0.165 |
| potassium | 1 | Clinical Chemistry | Physiology | -0.071 | -0.221 | 0.080 | 0.077 | -0.007 | -0.173 | 0.158 | 0.084 | 0.070 | 0.048 | 0.093 | 0.012 |
| pq | 1 | Electrocardiogram (ECG) | Heart | -0.065 | -0.154 | 0.024 | 0.045 | -0.065 | -0.127 | -0.003 | 0.032 | 0.002 | -0.026 | 0.029 | 0.014 |
| pr | 1 | Electrocardiogram (ECG) | Heart | -0.056 | -0.105 | -0.008 | 0.025 | -0.075 | -0.124 | -0.027 | 0.025 | -0.018 | -0.032 | -0.005 | 0.007 |
| qrs | 1 | Electrocardiogram (ECG) | Heart | 0.073 | 0.035 | 0.110 | 0.019 | 0.068 | 0.030 | 0.106 | 0.019 | -0.005 | -0.015 | 0.005 | 0.005 |
| qtc | 1 | Electrocardiogram (ECG) | Heart | 0.033 | -0.010 | 0.076 | 0.022 | 0.031 | -0.021 | 0.083 | 0.026 | -0.001 | -0.009 | 0.008 | 0.004 |
| qtc dispersion | 1 | Electrocardiogram (ECG) | Heart | 0.003 | -0.052 | 0.059 | 0.028 | -0.005 | -0.106 | 0.097 | 0.052 | -0.008 | -0.051 | 0.036 | 0.022 |
| red blood cell count | 1 | Hematology | Hematology | 0.077 | 0.007 | 0.147 | 0.036 | 0.100 | 0.032 | 0.168 | 0.035 | 0.023 | 0.009 | 0.037 | 0.007 |
| red blood cell distribution width | 1 | Hematology | Hematology | 0.125 | -0.004 | 0.253 | 0.065 | 0.135 | -0.004 | 0.274 | 0.071 | 0.010 | -0.003 | 0.024 | 0.007 |
| respiration rate | 1 | Echo | Heart | -0.138 | -0.218 | -0.059 | 0.041 | -0.070 | -0.180 | 0.039 | 0.056 | 0.061 | 0.023 | 0.099 | 0.020 |
| respiratory exchange ratio | 1 | Indirect Calorimetry | Metabolism | -0.012 | -0.090 | 0.066 | 0.040 | -0.011 | -0.088 | 0.067 | 0.039 | 0.002 | -0.006 | 0.009 | 0.004 |
| right anterior chamber depth | 1 | Eye Morphology | Eye | -0.449 | -1.329 | 0.431 | 0.449 | -0.416 | -1.292 | 0.460 | 0.447 | 0.032 | 0.026 | 0.037 | 0.003 |
| right corneal thickness | 1 | Eye Morphology | Eye | -0.036 | -0.228 | 0.157 | 0.098 | -0.031 | -0.196 | 0.135 | 0.085 | -0.001 | -0.024 | 0.021 | 0.011 |
| right inner nuclear layer | 1 | Eye Morphology | Eye | -0.255 | -0.763 | 0.254 | 0.260 | -0.279 | -0.837 | 0.280 | 0.285 | -0.018 | -0.066 | 0.031 | 0.025 |
| right outer nuclear layer | 1 | Eye Morphology | Eye | 0.006 | -0.078 | 0.090 | 0.043 | 0.011 | -0.073 | 0.095 | 0.043 | 0.006 | 0.000 | 0.011 | 0.003 |
| right posterior chamber depth | 1 | Eye Morphology | Eye | -0.078 | -0.291 | 0.135 | 0.109 | -0.076 | -0.289 | 0.136 | 0.109 | 0.007 | -0.018 | 0.032 | 0.013 |
| right total retinal thickness | 1 | Eye Morphology | Eye | -0.199 | -0.646 | 0.248 | 0.228 | -0.193 | -0.629 | 0.243 | 0.222 | 0.005 | -0.005 | 0.015 | 0.005 |
| rmssd | 1 | Electrocardiogram (ECG) | Heart | 0.180 | -0.088 | 0.448 | 0.137 | -0.016 | -0.411 | 0.379 | 0.202 | -0.118 | -0.245 | 0.009 | 0.065 |
| rp macrophage (cd19- cd11c-) | 1 | Immunophenotyping | Immunology | -0.077 | -0.340 | 0.187 | 0.134 | -0.075 | -0.335 | 0.186 | 0.133 | -0.075 | -0.207 | 0.058 | 0.068 |
| rr | 1 | Electrocardiogram (ECG) | Heart | -0.076 | -0.188 | 0.035 | 0.057 | -0.090 | -0.206 | 0.027 | 0.060 | -0.013 | -0.021 | -0.004 | 0.005 |
| sodium | 1 | Clinical Chemistry | Physiology | 0.026 | -0.117 | 0.170 | 0.073 | 0.034 | -0.134 | 0.201 | 0.085 | 0.010 | 0.007 | 0.013 | 0.002 |
| spleen weight | 1 | Immunophenotyping | Immunology | 0.187 | -0.050 | 0.425 | 0.121 | 0.113 | -0.160 | 0.387 | 0.140 | -0.154 | -0.210 | -0.098 | 0.029 |
| st | 1 | Electrocardiogram (ECG) | Heart | 0.003 | -0.054 | 0.061 | 0.029 | -0.005 | -0.081 | 0.070 | 0.039 | -0.003 | -0.018 | 0.011 | 0.007 |
| stroke volume | 1 | Echo | Heart | 0.059 | -0.078 | 0.197 | 0.070 | 0.157 | 0.009 | 0.306 | 0.076 | 0.094 | 0.078 | 0.110 | 0.008 |
| tibia length | 1 | Heart Weight | Morphology | -0.148 | -0.440 | 0.145 | 0.149 | -0.137 | -0.426 | 0.151 | 0.147 | 0.010 | 0.006 | 0.013 | 0.002 |
| total bilirubin | 1 | Clinical Chemistry | Physiology | 0.061 | -0.010 | 0.131 | 0.036 | 0.002 | -0.086 | 0.091 | 0.045 | -0.055 | -0.098 | -0.012 | 0.022 |
| total cholesterol | 1 | Clinical Chemistry | Physiology | 0.094 | -0.075 | 0.264 | 0.086 | 0.314 | 0.113 | 0.516 | 0.103 | 0.203 | 0.175 | 0.230 | 0.014 |
| total food intake | 1 | Indirect Calorimetry | Metabolism | -0.119 | -0.254 | 0.016 | 0.069 | -0.096 | -0.256 | 0.064 | 0.082 | 0.027 | -0.023 | 0.077 | 0.026 |
| total protein | 1 | Clinical Chemistry | Physiology | -0.042 | -0.062 | -0.022 | 0.010 | -0.036 | -0.062 | -0.009 | 0.013 | 0.009 | -0.001 | 0.019 | 0.005 |
| total water intake | 1 | Indirect Calorimetry | Metabolism | -0.146 | -0.237 | -0.054 | 0.047 | -0.210 | -0.268 | -0.151 | 0.030 | -0.065 | -0.137 | 0.007 | 0.037 |
| triglycerides | 1 | Clinical Chemistry | Physiology | -0.032 | -0.123 | 0.059 | 0.047 | 0.327 | 0.209 | 0.445 | 0.060 | 0.347 | 0.259 | 0.436 | 0.045 |
| urea (blood urea nitrogen - bun) | 1 | Clinical Chemistry | Physiology | -0.141 | -0.266 | -0.015 | 0.064 | -0.095 | -0.251 | 0.061 | 0.079 | 0.040 | 0.005 | 0.075 | 0.018 |
| uric acid | 1 | Clinical Chemistry | Physiology | 0.037 | -0.066 | 0.139 | 0.052 | 0.363 | 0.091 | 0.634 | 0.138 | 0.447 | -0.080 | 0.975 | 0.269 |
| white blood cell count | 1 | Hematology | Hematology | -0.091 | -0.170 | -0.011 | 0.041 | 0.117 | -0.002 | 0.236 | 0.061 | 0.198 | 0.137 | 0.259 | 0.031 |
| whole arena average speed | 1 | Open Field | Behaviour | -0.016 | -0.086 | 0.054 | 0.036 | -0.114 | -0.184 | -0.044 | 0.036 | -0.100 | -0.152 | -0.048 | 0.027 |
| whole arena resting time | 1 | Open Field | Behaviour | -0.053 | -0.101 | -0.005 | 0.025 | -0.059 | -0.108 | -0.011 | 0.025 | 0.005 | -0.051 | 0.061 | 0.029 |

#### 6.4 Table 6.4

```
# PDF pander(overall2, caption = 'Table 6.4: Summary of overall meta-analyses on
# the functional trait group level (GroupingTerm). Results for lnCVR, lnVR and
# lnRR and their respective upper and lower 95 percent CI's, standard error and
# I2 values are provided.')

# HTML
kable(overall2, digits = 3, caption = "Summary of overall meta-analyses on the functional trait group level (GroupingTerm). Results for lnCVR, lnVR and lnRR and their respective upper and lower 95 percent CI's, standard error and I2 values are provided.") %>% 
    kable_styling() %>% scroll_box(width = "100%", height = "200px")
```

Table 6.4: Summary of overall meta-analyses on the functional trait group level (GroupingTerm). Results for lnCVR, lnVR and lnRR and their respective upper and lower 95 percent CI’s, standard error and I2 values are provided.

| GroupingTerm | lnCVR | lnCVR\_lower | lnCVR\_upper | lnCVR\_se | lnCVR\_I2 | lnVR | lnVR\_lower | lnVR\_upper | lnVR\_se | lnVR\_I2 | lnRR | lnRR\_lower | lnRR\_upper | lnRR\_se | lnRR\_I2 |
| --- | --- | --- | --- | --- | --- | --- | --- | --- | --- | --- | --- | --- | --- | --- | --- |
| Behaviour | -0.004 | -0.024 | 0.017 | 0.010 | 39.033 | -0.018 | -0.074 | 0.038 | 0.029 | 92.799 | -0.020 | -0.063 | 0.024 | 0.022 | 89.238 |
| Morphology | 0.077 | 0.041 | 0.113 | 0.018 | 67.282 | 0.151 | 0.082 | 0.221 | 0.035 | 90.626 | 0.068 | 0.007 | 0.128 | 0.031 | 99.698 |
| Metabolism | -0.043 | -0.113 | 0.026 | 0.035 | 84.801 | 0.091 | -0.034 | 0.216 | 0.064 | 96.783 | 0.142 | 0.036 | 0.248 | 0.054 | 99.377 |
| Physiology | 0.013 | -0.014 | 0.039 | 0.014 | 70.566 | 0.036 | -0.028 | 0.100 | 0.033 | 90.643 | 0.016 | -0.044 | 0.077 | 0.031 | 99.817 |
| Immunology | -0.068 | -0.098 | -0.038 | 0.015 | 40.769 | -0.111 | -0.162 | -0.060 | 0.026 | 58.020 | -0.057 | -0.107 | -0.008 | 0.025 | 96.148 |
| Hematology | 0.022 | -0.017 | 0.060 | 0.020 | 60.753 | 0.080 | 0.032 | 0.129 | 0.025 | 64.943 | 0.039 | -0.002 | 0.080 | 0.021 | 99.630 |
| Heart | 0.018 | -0.013 | 0.050 | 0.016 | 81.298 | -0.005 | -0.036 | 0.026 | 0.016 | 74.058 | -0.005 | -0.032 | 0.023 | 0.014 | 98.985 |
| Hearing | 0.016 | -0.011 | 0.043 | 0.014 | 21.756 | 0.011 | -0.023 | 0.044 | 0.017 | 22.797 | -0.013 | -0.034 | 0.007 | 0.010 | 79.486 |
| Eye | -0.082 | -0.148 | -0.016 | 0.034 | 49.810 | -0.074 | -0.138 | -0.011 | 0.032 | 48.309 | 0.009 | 0.001 | 0.017 | 0.004 | 90.528 |
| All | 0.005 | -0.009 | 0.018 | 0.007 | 76.486 | 0.016 | -0.008 | 0.039 | 0.012 | 90.760 | 0.012 | -0.006 | 0.031 | 0.009 | 99.716 |

#### 6.5 Table 6.5

```
female <- (overall.female.plot3)
female$Sex <- c("female")

male <- (overall.male.plot3)
male$Sex <- c("male")

mf_sig <- rbind(female, male)
mf_sig <- mf_sig[, c(1, 17, 2:16)]

mf_sig <- mf_sig %>% arrange(desc(GroupingTerm))

# PDF pander(mf_sig, caption = 'Table 6.5: Provides an overview of meta-analysis
# results performed on traits that were significantly biased towards either sex.
# This table summarizes findings for both sexes and the respective functional
# trait groups.') HTML
kable(mf_sig, digits = 3, caption = "Provides an overview of meta-analysis results performed on traits that were significantly biased towards either sex. This table summarizes findings for both sexes and the respective functional trait groups.") %>% 
    kable_styling() %>% scroll_box(width = "100%", height = "300px")
```

Table 6.5: Provides an overview of meta-analysis results performed on traits that were significantly biased towards either sex. This table summarizes findings for both sexes and the respective functional trait groups.

| GroupingTerm | Sex | lnCVR | lnCVR\_lower | lnCVR\_upper | lnCVR\_se | lnCVR\_I2 | lnVR | lnVR\_lower | lnVR\_upper | lnVR\_se | lnVR\_I2 | lnRR | lnRR\_lower | lnRR\_upper | lnRR\_se | lnRR\_I2 |
| --- | --- | --- | --- | --- | --- | --- | --- | --- | --- | --- | --- | --- | --- | --- | --- | --- |
| Behaviour | female | -0.053 | -0.101 | -0.005 | 0.025 | 0.000 | -0.093 | -0.120 | -0.066 | 0.014 | 42.139 | -0.094 | -0.117 | -0.072 | 0.011 | 0.000 |
| Behaviour | male | 0.107 | 0.050 | 0.165 | 0.029 | 0.000 | 0.284 | 0.189 | 0.378 | 0.048 | 37.808 | 0.203 | 0.135 | 0.272 | 0.035 | 0.000 |
| Morphology | female | NA | NA | NA | NA | NA | NA | NA | NA | NA | NA | -0.110 | -0.230 | 0.011 | 0.061 | 93.935 |
| Morphology | male | 0.127 | 0.086 | 0.168 | 0.021 | 41.546 | 0.217 | 0.155 | 0.280 | 0.032 | 84.378 | 0.120 | 0.058 | 0.183 | 0.032 | 99.674 |
| Metabolism | female | -0.125 | -0.166 | -0.083 | 0.021 | 13.324 | -0.210 | -0.268 | -0.151 | 0.030 | 0.000 | NA | NA | NA | NA | NA |
| Metabolism | male | 0.094 | 0.052 | 0.136 | 0.021 | 0.000 | 0.224 | 0.100 | 0.348 | 0.063 | 95.761 | 0.223 | 0.110 | 0.335 | 0.058 | 98.647 |
| Physiology | female | -0.047 | -0.064 | -0.030 | 0.009 | 0.610 | -0.163 | -0.296 | -0.030 | 0.068 | 92.843 | -0.117 | -0.221 | -0.014 | 0.053 | 99.301 |
| Physiology | male | 0.096 | 0.063 | 0.130 | 0.017 | 0.109 | 0.240 | 0.159 | 0.320 | 0.041 | 56.006 | 0.145 | 0.071 | 0.218 | 0.038 | 98.518 |
| Immunology | female | -0.090 | -0.148 | -0.032 | 0.030 | 81.548 | -0.180 | -0.218 | -0.141 | 0.020 | 0.000 | -0.159 | -0.229 | -0.090 | 0.035 | 94.729 |
| Immunology | male | NA | NA | NA | NA | NA | NA | NA | NA | NA | NA | 0.028 | 0.001 | 0.055 | 0.014 | 44.587 |
| Hematology | female | -0.092 | -0.150 | -0.034 | 0.030 | 0.000 | NA | NA | NA | NA | NA | -0.013 | -0.018 | -0.007 | 0.003 | 84.990 |
| Hematology | male | 0.082 | 0.048 | 0.117 | 0.017 | 0.089 | 0.144 | 0.046 | 0.243 | 0.050 | 86.350 | 0.116 | 0.027 | 0.205 | 0.046 | 98.445 |
| Heart | female | -0.091 | -0.171 | -0.012 | 0.041 | 66.548 | -0.084 | -0.109 | -0.059 | 0.013 | 0.528 | -0.108 | -0.179 | -0.036 | 0.036 | 98.814 |
| Heart | male | 0.154 | 0.085 | 0.223 | 0.035 | 78.699 | 0.104 | 0.062 | 0.146 | 0.022 | 36.412 | 0.058 | 0.041 | 0.075 | 0.009 | 84.650 |
| Hearing | female | NA | NA | NA | NA | NA | NA | NA | NA | NA | NA | -0.035 | -0.062 | -0.009 | 0.014 | 61.362 |
| Hearing | male | NA | NA | NA | NA | NA | 0.087 | 0.007 | 0.167 | 0.041 | 0.000 | 0.018 | 0.006 | 0.031 | 0.006 | 0.000 |
| Eye | female | -0.164 | -0.250 | -0.078 | 0.044 | 3.071 | -0.166 | -0.277 | -0.056 | 0.057 | 24.076 | NA | NA | NA | NA | NA |
| Eye | male | NA | NA | NA | NA | NA | NA | NA | NA | NA | NA | 0.019 | 0.004 | 0.034 | 0.008 | 97.046 |
| All | female | -0.087 | -0.109 | -0.064 | 0.012 | 66.479 | -0.130 | -0.157 | -0.103 | 0.014 | 75.919 | -0.098 | -0.129 | -0.068 | 0.016 | 99.612 |
| All | male | 0.111 | 0.093 | 0.130 | 0.009 | 34.070 | 0.198 | 0.162 | 0.233 | 0.018 | 86.497 | 0.111 | 0.083 | 0.138 | 0.014 | 99.639 |

#### 6.6 Table 6.6

```
het <- heterog6 %>% pivot_wider(names_from = parameter, values_from = value:ci.h) %>% 
    rename(lnCVR = value_heteroCVR, lnCVR_lower = ci.low_heteroCVR, lnCVR_higher = ci.high_heteroCVR, 
        lnVR = value_heteroVR, lnVR_lower = ci.low_heteroVR, lnVR_higher = ci.high_heteroVR, 
        lnRR = value_heteroRR, lnRR_lower = ci.low_heteroRR, lnRR_higher = ci.high_heteroRR, 
        CVR = mean_heteroCVR, VR = mean_heteroVR, RR = mean_heteroRR, CVR_lower = ci.l_heteroCVR, 
        VR_lower = ci.l_heteroVR, RR_lower = ci.l_heteroRR, CVR_higher = ci.h_heteroCVR, 
        VR_higher = ci.h_heteroVR, RR_higher = ci.h_heteroRR) %>% select(1, 2, 5, 
    8, 3, 6, 9, 4, 7, 10, 14, 17, 20, 15, 18, 21, 16, 19, 22)

# PDF pander(het, caption = 'Table 6.6: Summarizes our findings on heterogeneity
# due to institutions and mouse strains. These results are based on meta-analyses
# on sigma^2 and errors for mouse strains and centers (Institutions), following
# the identical workflow from above.')

# HTML
kable(het, digits = 3, caption = "Summarizes our findings on heterogeneity due to institutions and mouse strains. These results are based on meta-analyses on sigma^2 and errors for mouse strains and centers (Institutions), following the identical workflow from above.") %>% 
    kable_styling() %>% scroll_box(width = "100%", height = "200px")
```

Table 6.6: Summarizes our findings on heterogeneity due to institutions and mouse strains. These results are based on meta-analyses on sigma^2 and errors for mouse strains and centers (Institutions), following the identical workflow from above.

| GroupingTerm | lnCVR | lnCVR\_lower | lnCVR\_higher | lnVR | lnVR\_lower | lnVR\_higher | lnRR | lnRR\_lower | lnRR\_higher | CVR | CVR\_lower | CVR\_higher | VR | VR\_lower | VR\_higher | RR | RR\_lower | RR\_higher |
| --- | --- | --- | --- | --- | --- | --- | --- | --- | --- | --- | --- | --- | --- | --- | --- | --- | --- | --- |
| Behaviour | -1.649 | -2.464 | -0.835 | -1.545 | -2.359 | -0.730 | -2.640 | -4.135 | -1.145 | 0.192 | 0.085 | 0.434 | 0.213 | 0.094 | 0.482 | 0.071 | 0.016 | 0.318 |
| Morphology | -1.745 | -2.424 | -1.066 | -1.725 | -2.404 | -1.046 | -7.277 | -10.004 | -4.549 | 0.175 | 0.089 | 0.344 | 0.178 | 0.090 | 0.351 | 0.001 | 0.000 | 0.011 |
| Metabolism | -2.404 | -3.567 | -1.240 | -2.897 | -4.260 | -1.534 | -2.970 | -4.134 | -1.806 | 0.090 | 0.028 | 0.289 | 0.055 | 0.014 | 0.216 | 0.051 | 0.016 | 0.164 |
| Physiology | -2.899 | -3.762 | -2.037 | -1.922 | -2.688 | -1.155 | -2.664 | -3.430 | -1.898 | 0.055 | 0.023 | 0.130 | 0.146 | 0.068 | 0.315 | 0.070 | 0.032 | 0.150 |
| Immunology | -0.291 | -0.795 | 0.213 | -0.231 | -0.735 | 0.273 | -0.895 | -1.790 | 0.001 | 0.748 | 0.452 | 1.238 | 0.793 | 0.479 | 1.313 | 0.409 | 0.167 | 1.001 |
| Hematology | -0.463 | -1.120 | 0.193 | -0.369 | -1.026 | 0.287 | -1.287 | -2.220 | -0.354 | 0.629 | 0.326 | 1.213 | 0.691 | 0.359 | 1.333 | 0.276 | 0.109 | 0.702 |
| Heart | -3.573 | -4.272 | -2.874 | -2.980 | -3.549 | -2.411 | -3.877 | -4.443 | -3.311 | 0.028 | 0.014 | 0.056 | 0.051 | 0.029 | 0.090 | 0.021 | 0.012 | 0.036 |
| Hearing | -3.059 | -5.049 | -1.068 | -2.361 | -4.352 | -0.370 | -3.602 | -5.592 | -1.611 | 0.047 | 0.006 | 0.344 | 0.094 | 0.013 | 0.690 | 0.027 | 0.004 | 0.200 |
| Eye | -3.707 | -5.102 | -2.312 | -3.228 | -4.331 | -2.125 | -5.773 | -6.574 | -4.972 | 0.025 | 0.006 | 0.099 | 0.040 | 0.013 | 0.119 | 0.003 | 0.001 | 0.007 |
| All | -2.290 | -2.648 | -1.932 | -1.936 | -2.254 | -1.619 | -3.452 | -3.944 | -2.960 | 0.101 | 0.071 | 0.145 | 0.144 | 0.105 | 0.198 | 0.032 | 0.019 | 0.052 |

### 7 R Session Information

```
sessionInfo() %>% pander()
```

**R version 3.6.0 (2019-04-26)**

**Platform:** x86\_64-apple-darwin15.6.0 (64-bit)

**locale:**
en\_AU.UTF-8||en\_AU.UTF-8||en\_AU.UTF-8||C||en\_AU.UTF-8||en\_AU.UTF-8

**attached base packages:**
*grid*, *stats*, *graphics*, *grDevices*, *utils*, *datasets*, *methods* and *base*

**other attached packages:**
*splus2R(v.1.2-2)*, *pander(v.0.6.3)*, *knitr(v.1.29)*, *here(v.0.1)*, *png(v.0.1-7)*, *ggpubr(v.0.2.1)*, *magrittr(v.1.5)*, *robumeta(v.2.0)*, *kableExtra(v.1.1.0)*, *forcats(v.0.4.0)*, *stringr(v.1.4.0)*, *tidyr(v.1.0.2)*, *tibble(v.3.0.3)*, *ggplot2(v.3.3.2)*, *tidyverse(v.1.2.1)*, *purrr(v.0.3.2)*, *devtools(v.2.1.0)*, *usethis(v.1.5.1)*, *metafor(v.2.1-0)*, *Matrix(v.1.2-17)*, *dplyr(v.0.8.3)*, *readr(v.1.3.1)* and *pacman(v.0.5.1)*

**loaded via a namespace (and not attached):**
*nlme(v.3.1-140)*, *fs(v.1.3.1)*, *lubridate(v.1.7.4)*, *RColorBrewer(v.1.1-2)*, *webshot(v.0.5.1)*, *httr(v.1.4.0)*, *rprojroot(v.1.3-2)*, *tools(v.3.6.0)*, *backports(v.1.1.4)*, *utf8(v.1.1.4)*, *R6(v.2.4.0)*, *mgcv(v.1.8-28)*, *colorspace(v.1.4-1)*, *withr(v.2.1.2)*, *tidyselect(v.1.1.0)*, *prettyunits(v.1.0.2)*, *processx(v.3.4.0)*, *compiler(v.3.6.0)*, *cli(v.1.1.0)*, *rvest(v.0.3.4)*, *formatR(v.1.7)*, *xml2(v.1.2.1)*, *desc(v.1.2.0)*, *labeling(v.0.3)*, *bookdown(v.0.20)*, *scales(v.1.1.1)*, *callr(v.3.3.0)*, *digest(v.0.6.20)*, *rmarkdown(v.2.3)*, *pkgconfig(v.2.0.2)*, *htmltools(v.0.3.6)*, *sessioninfo(v.1.1.1)*, *highr(v.0.8)*, *rlang(v.0.4.7)*, *readxl(v.1.3.1)*, *rstudioapi(v.0.10)*, *farver(v.2.0.3)*, *generics(v.0.0.2)*, *jsonlite(v.1.6)*, *fansi(v.0.4.0)*, *Rcpp(v.1.0.1)*, *munsell(v.0.5.0)*, *lifecycle(v.0.2.0)*, *stringi(v.1.4.3)*, *yaml(v.2.2.0)*, *pkgbuild(v.1.0.3)*, *crayon(v.1.3.4)*, *lattice(v.0.20-38)*, *haven(v.2.1.1)*, *cowplot(v.1.0.0)*, *splines(v.3.6.0)*, *hms(v.0.5.0)*, *ps(v.1.3.0)*, *pillar(v.1.4.6)*, *ggsignif(v.0.5.0)*, *codetools(v.0.2-16)*, *pkgload(v.1.0.2)*, *glue(v.1.3.1)*, *evaluate(v.0.14)*, *remotes(v.2.1.0)*, *modelr(v.0.1.4)*, *vctrs(v.0.3.2)*, *testthat(v.2.1.1)*, *cellranger(v.1.1.0)*, *gtable(v.0.3.0)*, *assertthat(v.0.2.1)*, *xfun(v.0.16)*, *broom(v.0.7.0)*, *viridisLite(v.0.3.0)*, *memoise(v.1.1.0)* and *ellipsis(v.0.3.1)*
